# Structural and functional clues challenge the hypothesis that the *yjdF* riboswitch is natively regulated through broad recognition of azaaromatic compounds

**DOI:** 10.1101/2025.06.18.660367

**Authors:** Savannah F. Spradlin, Kyle A. Dickerson, Robert T. Batey

## Abstract

While most riboswitches are highly selective for their cognate ligand, the *yjdF* riboswitch is distinct in its ability to bind a broad set of aromatic compounds. This observation has led to the hypothesis that this RNA is regulated by toxic azaaromatic compounds, triggering a detoxification mechanism by activating translation of the YjdF protein in response to ligand binding. To understand how these compounds turn on gene expression by the *yjdF* riboswitch, we determined the crystal structure of the *Bacillus subtilis yjdF* riboswitch in complex with activating (chelerythrine) and non-activating (lumichrome) ligands. These structures reveal that the RNA binds these compounds in a near-identical fashion, adopting the same local and global conformation. However, the unexpected extension of the regulatory helix through formation of several base pairs from highly conserved nucleotides suggests that this element plays an important role in ligand-dependent gene expression. Using a reporter assay in *B. subtilis*, we found that chelerythrine-dependent activation is insensitive to mutation of these conserved nucleotides that are essential for activation of the riboswitch. These data suggest that the *yjdF* riboswitch is responsive to a yet unknown cellular metabolite and remains an orphan riboswitch.

**GRAPHICAL ABSTRACT:** 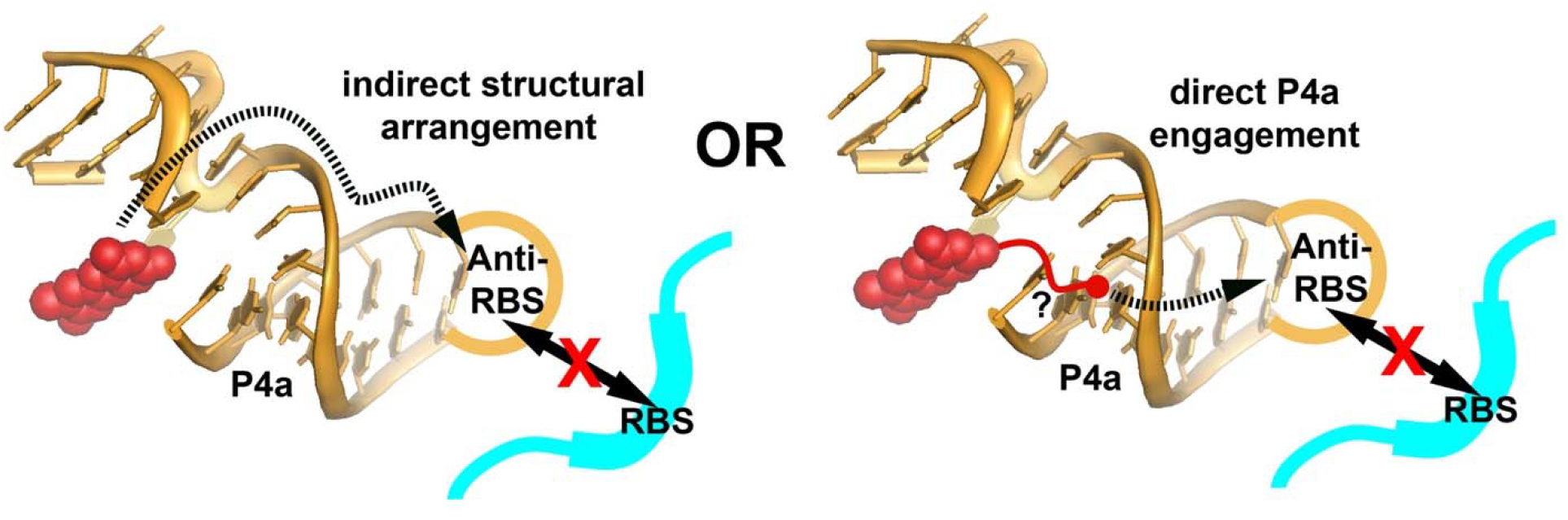

## Introduction

Riboswitches are regulatory elements typically found in the 5’-leader sequence of bacterial mRNAs that govern expression through their ability to directly interact with small molecules or ions (1,2). The typical riboswitch comprises two functional domains: the ligand-binding aptamer domain and a downstream regulatory switch called the expression platform (3). The ligand occupancy status of the aptamer domain is read out by the expression platform, which adopts one of two mutually exclusive conformational states: one enabling expression of the downstream gene (ON state) or one that represses the expression of the gene (OFF state). This mechanism of gene regulation is common in bacteria, particularly in Firmicutes and Fusobacteria, where these regulatory elements control expression of a broad array of genes involved in normal cellular homeostasis, second messenger signaling, and virulence (4,5).

Currently, over 55 riboswitches have been identified and validated (6), each binding a specific small molecule using a distinct secondary and tertiary architecture (7,8). Almost invariably, these aptamers are highly specific for their cognate ligand and discriminate against closely related compounds in the metabolome. For example, the lysine riboswitch has a large binding pocket that extensively recognizes the amino acid’s main chain atoms through hydrogen bonding interactions and the epsilon-amino group through an electrostatic interaction (9,10). Despite the fact that many of the proteogenic amino acids can sterically fit into the binding pocket, the riboswitch discriminates between lysine and serine by at least 10,000-fold in affinity (11). Similarly, *S*-adenosylmethionine (SAM) binding riboswitches are capable of up to 500-fold discrimination between SAM and its demethylated product, *S*-adenosylhomocysteine (SAH) (12,13). Other riboswitches, such as the tetrahydrofolate (THF) (14,15) and the nicotinamide adenine dinucleotide (NAD^+^)-II (16) riboswitches discriminate between the oxidized and reduced forms of their cognate metabolite. Thus, many riboswitches achieve levels of molecular discrimination that rival their protein counterparts, enabling them to regulate gene expression by sensing the intracellular concentration of a single metabolite or a small set of related metabolites.

A striking and unique counterexample to the high selectivity of typical aptamer domains was the discovery of a riboswitch that controls *yjdF* genes found primarily in Firmicutes and Actinobacteria (17). The cognate ligand of a riboswitch aptamer domain is generally identified using knowledge of the downstream genes that they regulate (18,19). However, the *yjdF* riboswitch motif is found upstream almost exclusively of the YjdF protein, whose function is currently unknown (17). To determine the cognate ligand, the Breaker group conducted an exhaustive survey of compounds with the *Bacillus subtilis (Bsu) yjdF* aptamer domain *in vitro* using in-line probing and an *in vivo* reporter assay using the full riboswitch in *B. subtilis* (17). Their analysis revealed that *yjdF* interacts with a broad array of compounds, unlike other riboswitches. The compounds all share the commonality of containing at least two heterocyclic aromatic (azaaromatic) rings (*Figure 1A*), leading to the conclusion that this riboswitch recognizes compounds with degenerate specificity—that is, recognizing a set of compounds that share common structural or chemical features but are not closely related (20-22). Despite no clear common chemical structure shared by *yjdF*-binding compounds beyond an azaaromatic ring system, a number of these compounds bound with low-to-mid nanomolar affinities (17).

**Figure 1.**
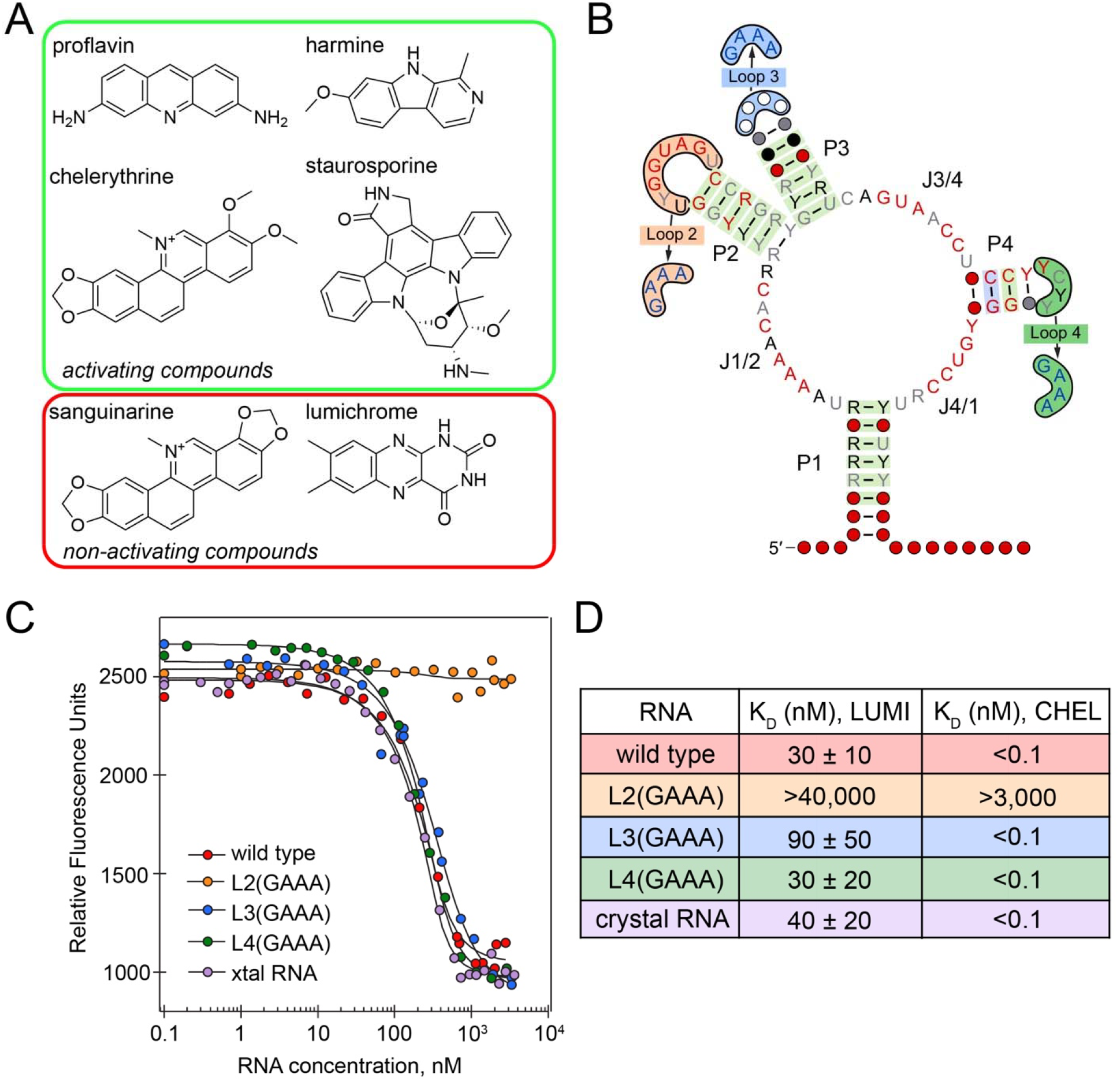
Ligand binding and secondary structural features of the *yjdF* riboswitch aptamer domain. (A) Chemical structures of compounds identified as translational activators and non-activators by the Breaker group (17). All the compounds depicted have low-to-mid nanomolar binding affinities for the *B. subtilis yjdF* riboswitch. (B) Secondary structure derived from covariation analysis of 507 *yjdF* sequences available in the Rfam 15.0 database (42). Nucleotide identity and base pairs are denoted and colored according to the standard R2R convention (44). Loops mutated to GAAA tetraloops to test their effect on azaaromatic compound binding are noted. (C) Representative binding curves for the wild type and select mutant *B. subtilis yjdF* riboswitch aptamer domains binding to lumichrome. (D) Binding data for lumichrome and chelerythrine (150 mM KCl and 5 mM Mg^2+^). Note that under these conditions, the affinity of chelerythrine for wild type RNA was so high that the K_D_ could not be accurately determined (41) and could only yield a determination that it is below 100 pM.

While a broad set of azaaromatic compounds bind the *yjdF* riboswitch aptamer domain, not all binders are able to switch genetic regulation (17). The *yjdF* riboswitch is always observed as an “ON” switch such that effector binding prevents the aptamer domain from interacting with the ribosome binding site (RBS), enabling the ribosome to translate the message. The terminal loop L4 of the aptamer domain (**Figure 1B**) contains a sequence complementary to the RBS that base pairs and blocks translation in the absence of an effector. A cell-based reporter assay demonstrated that only a subset of compounds that bind the aptamer derepress expression of a reporter gene (17). Notably, structurally similar compounds such as chelerythrine and sanguinarine (**Figure 1A**) differed in their ability to regulate gene expression despite both binding the riboswitch with low nanomolar affinity, confounding a clear understanding of the specificity of this riboswitch. However, this phenomenon is not unique to the *yjdF* riboswitch. For example, the THF-I riboswitch binds compounds such as 2,6-diaminopurine with higher affinity than THF, but some of these high affinity ligands only modestly regulate expression resulting is a disconnect between RNA binding and regulatory activity (23). Thus, for *yjdF*, the chemical features of the effector required for ligand binding and promoting gene regulation may be distinct from each other.

Amongst the compounds surveyed by the Breaker group using an *in vivo* reporter assay, there were no known natural bacterial metabolites that activated expression by the *yjdF* riboswitch and only five aromatic compounds (17). These results, alongside their *in vitro* binding analysis led to the hypothesis that this riboswitch responds broadly to toxic azaaromatic compounds, and that the YjdF protein is likely involved in attenuating the effect of these compounds. Thus, the *yjdF* riboswitch was proposed to be analogous to the bacterial PadR regulatory protein that binds a broad array of planar hydrophobic compounds as part of a detoxification mechanism (24,25).

To understand the basis for promiscuous binding of azaaromatic compounds by the riboswitch and the effector molecule’s relationship to gene regulation, we determined the crystal structure of the *B. subtilis yjdF* riboswitch aptamer domain in complex with an activating chelerythrine, **Figure 1A**) and a non-activating (lumichrome, **Figure 1A**) compound. These structures reveal a T-shaped RNA in which the ligand binding site is comprised of a terminal loop motif docked into a four-way junction. The azaaromatic binding site is large and planar with one edge open to bulk solvent, revealing how the riboswitch accommodates ligands of diverse sizes and shapes. This is further reinforced by our finding that the *yjdF* aptamer can bind a broad array of fluorogenic dyes and turn on their fluorescence. However, both chelerythrine and lumichrome interact with the RNA in near-identical fashion, suggesting an unappreciated mode of linking effector binding to gene regulation. Using structure-guided mutagenesis and a cell-based reporter assay in *B. subtilis*, we investigated a region of the aptamer containing highly conserved nucleotides at the base of a stem-loop containing the anti-RBS sequence that may serve as a critical link between ligand binding and regulatory activity. These data suggest that formation of conserved pyrimidine-pyrimidine pairs is crucial for gene regulation but how effector binding is coupled to stabilization of these pairs remains unknown. Our structural and functional observations suggest that the identified planar aromatic compounds may not be the natural effector of the *yjdF* riboswitch, but rather a yet unidentified compound that directly interacts with the aromatic site and the conserved pyrimidine pairs.

## Materials and Methods

### RNA preparation

All RNAs were transcribed and purified using a standard *in vitro* T7 RNA polymerase (RNAP) reaction and purified by denaturing polyacrylamide gel electrophoresis (PAGE) (26,27). In brief, chemically synthesized ssDNA oligonucleotides (Integrated DNA Technologies) were assembled using recursive PCR (28) to create a dsDNA to be used as T7 RNAP transcription templates. For transcription templates used to synthesize RNAs for crystallography, the 5’ end of the outer 3’ PCR primer contained two 2’-*O*-methoxy groups to inhibit non-templated addition of nucleotides at the 3’ end of the product RNA (29). RNA was transcribed in a reaction containing T7 transcription buffer (40 mM Tris-HCl pH 8.0, 10 mM dithiothreitol (DTT), 2 mM spermidine, 0.01% Triton X-100), 24 mM MgCl_2_, 4 mM each NTP, 8 mM DTT, ∼1 µM dsDNA template, 4 units of inorganic pyrophosphatase, 0.5 mg/mL T7 of RNA polymerase for 2 hours at 37 °C. Product RNA was precipitated with ethanol and purified by electrophoresis through a 12% denaturing (8 M urea) 29:1 acrylamide:bisacrylamide polyacrylamide gel. RNA was imaged by UV shadowing and the product band was excised from the gel and eluted via passive elution overnight in ddH_2_O. The supernatant was buffer exchanged into 0.5x T.E. buffer (5 mM Tris-HCl, pH 8.0, 0.5 mM ethylenediaminetetraacetic acid (EDTA)) and concentrated to ∼0.5 mM using a 10 kDa MWCO Amicon® Ultra Centrifugal Filter and stored at -20 °C until use. Molar extinction coefficients for calculation of RNA concentration by their absorption at 260 nm were calculated using IDT Tools (idtdna.com). All RNA sequences used in this study are given in **Supplementary Table S1**.

### Crystallization

RNA was prepared for crystallization by switching into exchange buffer (20 mM Tris-HCl pH 8.0, 25 mM KCl, and 0.5 mM MgCl_2_). Solid ligand was dissolved in exchange buffer and filtered using a Corning® Costar® Spin-X® 0.22 μm cellulose acetate centrifuge tube filter. 200 μL of the ligand solution was added to the 200 µL of 450 µM RNA in a 0.5 mL 10 kDa MWCO Amicon® Ultra Centrifugal Filter, mixed by pipetting, and the total volume was reduced to 200 µL. This process was repeated until the ligand concentration in the flow through was the same as the input to ensure saturation of the RNA with ligand. RNA was crystallized using the hanging drop vapor diffusion method. 2 μL of the RNA-ligand mixture (approximately 500 µM lumichrome and 450 µM RNA) in exchange buffer was added to 1 or 2 μL mother liquor and equilibrated over 500 μL of 35% 2-methyl-2,4-pentanediol (MPD). For native lumichrome crystals, the mother liquor contained 6% MPD, 9 mM spermine, 40 mM Na-cacodylate pH 7.0, 20 mM KCl. Crystals used for phasing contained a mother liquor of 5% MPD, 3 mM spermine, 50 mM Na-cacodylate pH 7.0, 20 mM NaCl. The drops were equilibrated at 21 °C and crystals began to form between 3 and 7 days of equilibration. Native crystals were harvested from drops using a nylon loop, and frozen by immediately plunging into liquid nitrogen. To derivatize crystals with iridium, 1 µL of 10 mM iridium(III) hexammine was added directly to the drop containing the crystals and allowed to soak for 7.5 hours (30). Crystals were removed from the drop using a nylon loop and plunged immediately into liquid nitrogen. Crystals with chelerythrine were grown using a mother liquor containing 5% MPD, 3 mM spermine, 40 mM Na-cacodylate pH 7.0, 20 mM KCl, 5 mM BaCl_2_ and handled in the same fashion as lumichrome containing crystals.

### Structure determination and refinement

X-ray diffraction data for iridium-soaked crystals of the lumichrome-*yjdF* complex were collected at the Advanced Light Source at Lawrence Berkeley National Laboratory using beamline 8.2.1 and native datasets were acquired at the Macromolecular X-ray Crystallography Core at the University of Colorado Boulder with a RIGAKU MicroMax-003 X-ray source with a Dectris PILATUS 200K 2D hybrid pixel array detector. All datasets were integrated and scaled using HKL-3000 (31). Initial phases were calculated using data from iridium-soaked lumichrome-*yjdF* crystals using the AutoSol (32) to find 21 heavy atom sites and to calculate experimental phases in PHENIX (33) (all crystallographic data and refinement statistics are given in **Supplementary Table S2**). Initial building of an RNA model into a density modified experimental map was performed using iterative rounds of building and refinement in the AutoBuild wizard in PHENIX (34), resulting in most of the RNA modeled except for J3/4. Further iterative manual model building was performed in Coot (35) and refined in PHENIX. Following complete building of the RNA, lumichrome was placed into the model using LigandFit implemented in PHENIX (36). The local geometry of the structure was improved using ERRASER (37,38) prior to the last round of refinement in PHENIX.

A native lumichrome-*yjdF* model was generated using a home-source dataset of higher resolution and data quality collected on native crystals not soaked with iridium. Following integration and scaling of the native dataset using HKL-3000, the model from the iridium dataset was used in molecular replacement using Phaser (39). To reduce model bias, the test sets for the iridium and native datasets were matched and high-temperature simulated annealing was implemented in PHENIX during refinement (40). Iterative manual adjustment of the RNA in Coot and refinement in PHENIX were performed, followed by automated water picking. Metal ions were identified and placed using stereochemical considerations such as short M^2+^-oxygen bond distances (2.1 – 2.2 Å) and local electrostatic environment around the ion. Prior to final rounds of refinement, the local geometry of the structure was improved with two rounds of ERRASER. The final lumichrome-*yjdF* model was refined to a R_work_ and R_free_ of 0.216 and 0.247, respectively; representative simulated annealing omit and 2F_o_-F_c_ maps are shown in **Supplemental Figures S1 and S2**. The RNA from the native lumichrome-*yjdF* model was used as a search model in Phaser in PHENIX to calculate an electron density map for data collected from chelerythrine-*yjdF* crystals. Regions in L2, J3/4 and J1/2 were rebuilt and the ligand placed, resulting in a model with a final R_work_ and R_free_ of 0.230 and 0.263, respectively. 2F_o_-F_c_ and simulated annealing electron density maps of bound chelerythrine and adjacent nucleotides is shown in **Supplemental Figure S3**. Atomic coordinates and associated structure factors are deposited in the RSCB Protein Data Bank. Note that all nucleotide numbering in this study aligns with the *B. subtilis yjdF* RNA used for crystallography.

### Fluorescence measurements

Binding assays were guided by the framework described in Jarmoskaite *et al*. (41). To prepare RNA for binding experiments, RNA and Milli-Q water were added to a PCR tube and heated for two minutes at 95 °C followed by immediate incubation in a slushy ice bath to cool for at least 10 minutes. Then, an equal volume of room temperature 2x binding buffer was added and mixed by gentle pipetting; this stock of RNA was serially diluted for binding reactions. Three different binding buffers were used in this study (Buffer A: 20 mM Tris-HCl, pH 7.5, 150 mM KCl, 5 mM MgCl_2_, 0.001% Nonidet P-40; Buffer B: 20 mM Tris-HCl, pH 7.5, 50 mM KCl, 20 µM MgCl_2_, 0.001% Nonidet P-40; Buffer C: 20 mM Tris-HCl, pH 7.5, 135 mM KCl, 15 mM NaCl, 1 mM MgCl_2_) that differ in their salt concentrations. Ligands were dissolved in 1x binding buffer and diluted with 1x binding buffer. Binding reactions comprising 25 µL of ligand and 25 µL of serially diluted RNA were added to a Corning® Black polystyrene 384-Well Assay Plates (Flat Bottom, Low Flange, non-binding surface).

Fluorescence measurements were taken using a BMG Labtech CLARIOstar Plus plate reader. Prior to each experiment, plates were calibrated using the CLARIOstar plate mapping feature with 50 µL of 500 nM fluorescein in ddH_2_O. Focal heights were determined experimentally using the focal height adjustment. Gain values were kept constant between experiments. Optimal excitation and emission wavelengths were determined experimentally using excitation and emission scans as well as running multichromatic experiments and comparing their curve fits (**Supplementary Table S3**). Data generated was viewed in BMG MARS Data Analysis Software, then exported to Microsoft Excel which was used for data export, storage, technical replicate averaging, and all normalization. All data was acquired with a minimum of three technical replicates.

All data was analyzed using KaleidaGraph (Synergy Software, v 5.0.1). Curve fitting for lumichrome was done with the quadratic equation

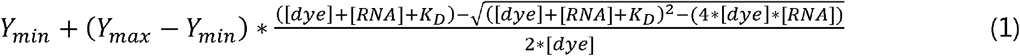

where Y_min_ is the lower baseline, Y_max_ is the upper baseline, [dye] is the concentration of fluorophore used for the experiment, [RNA] is the concentration of RNA, and K_D_ is the dissociation constant. For lumichrome, the dye concentration was spectroscopically determined and fixed at 380 nM for fitting and the chelerythrine concentrations were fixed at 0.5 nM.

Fluorescence fold turn-on measurements were taken at the excitation and emission wavelength and bandwidths specified in **Supplementary Table S3**. Fold turn-on calculations used two measurements: dye with RNA (20 µL of 1 µM fluorophore and 10 µM wild type *B. subtilis yjdF* RNA in 1x Buffer C) and dye alone (20 µL of 1 µM fluorophore in 1x Buffer C). The observed fold turn-on was determined by dividing the relative fluorescence in the presence of RNA by the relative fluorescence in the absence of RNA.

### Bioinformatic Analysis

All sequences used in bioinformatic analysis were downloaded from the Rfam (v. 15.0) database for *yjdF* RNA (RF01764) (42). We used the 507 *yjdF* RNA sequences and covariation file from Rfam to create a Stockholm alignment using Infernal (43) installed on a Windows Subsystem for Linux. This alignment, along with information about base-base association patterns observed within our crystal structures, was used to perform a structure-based analysis of conserved sequence-space. R2R (44) was used to generate a visualization of the generated Infernal alignment, and this secondary sequence and conservation pattern agrees with that previously derived by the Breaker lab (17).

To derive taxonomic information for the sequences in the alignment, we utilized a combination of automated scripts and code generated with guidance from ChatGPT run as Python scripts on Visual Studio Code. Accession IDs were processed through the National Center for Biotechnology Information (NCBI) Entrez Programming Utilities (E-utilities) to obtain corresponding taxonomic IDs. The resulting taxonomic IDs were then saved alongside their respective accession ID for verification and merging. To facilitate base conservation analysis, each nucleotide position was iteratively separated and numbered, and those reference position numbers were used to call base conservation and identity information. To assess conservation in the context of base pairs and triplets, a list was created with the reference position numbers of base pairs/triplets. A table was created containing all possible base pair/triplet identities, including deletions, which was then populated by those base pair occurrences. The *yjdF* alignment in FASTA format is given in **Supplementary Dataset 1**.

### Cell-based assays in Bacillus subtilis

A *yjdF-lacZ* reporter was constructed in *B. subtilis* strain *PY79* (45). For integration of the reporter into the *amyE* locus, an integration vector derived from pDR110 (ECE311, Bacillus Genetic Stock Center) was created using standard molecular biological techniques. The *Bsu yjdF* riboswitch DNA sequence (range 1277151 to 1277283, GenBank accession CP136402.1) was fused to a downstream *lacZ* reporter using recombinant PCR and the resultant gene inserted between the HindIII and SphI restriction sites of the multiple cloning site under transcriptional control of the IPTG-inducible *Pspank* promoter in pDR110. The resultant plasmid was transformed into *B. subtilis PY79* via natural competence followed by selection for spectinomycin resistance (46,47). Successful genomic integration of the construct at the *amyE* locus was validated by PCR amplification using *amyE* primers flanking the expected integration site and Nanopore sequencing of the amplicon. Strains containing mutants of the *Bsu yjdF* riboswitch were constructed and sequence verified in the same manner. Visualization of reporter activity was performed using an X-gal (5-bromo-4-chloro-3-indolyl-β-D-galactopyranoside) plate assay by streaking *yjdF* reporter wild type or mutant strains onto Chemical Salts Broth (CSB)-agar plates (1x CSB (48), 100 µg/mL spectinomycin, 1 mM IPTG, and 80 µg/mL X-gal) and CSB reporter plates containing 50 nM chelerythrine.

To quantify the activation of wild type and mutant *yjdF* riboswitches by chelerythrine, a standard Miller assay was performed (49). For each assay, *yjdF-lacZ* reporter strains of *Bacillus subtilis PY79* was grown overnight in 1 mL liquid CSB media supplemented with 100 µg/mL spectinomycin in a 10 mL capped plastic culture tube, inoculated with a single colony, and incubated at 37 °C in a rotating drum. The following morning, a fresh 10 mL CSB culture was inoculated using 1:100 of the overnight culture and grown in the same manner until the absorbance at 600 nm reached 0.4-0.5. This culture was then split into 3 separate 2 mL cultures: negative control, 1 mM IPTG and 1 mM IPTG plus 50 nM chelerythrine, 5 µM lumichrome, or 50 µM lumichrome and incubated at 37 °C. After 30 minutes, 1 mL of each culture condition was pipetted into two 1.5 mL microcentrifuge tubes and centrifuged in a tabletop centrifuge for 1 minute at 16,000 x*g* in a microcentrifuge. Cell pellets were each resuspended in 750 µL of working buffer (60 mM Na_2_HPO4, 40 mM NaH_2_PO_4_, 10 mM KCl, 1 mM MgCl_2_ and 20 mM β-mercaptoethanol) and combined. This cell pellet resuspension was then divided into three separate reactions of 500 µL as technical replicates. Using a blank of working buffer, the absorbance at 600 nm of each 500 µL reaction were measured. Each reaction was lysed by adding 6.25 µL of 15 mg/mL lysozyme, briefly vortexing, then incubating at 37 °C for 20 minutes. Then, upon adding 93.8 µL of 4 mg/mL *ortho*-nitrophenyl-β-D-galactopyranoside (ONPG) the start time was recorded and the reaction mixed well by pipetting. 15 minutes after the reaction start time, the reaction was quenched using 250 µL of 1 M Na_2_CO_3_. Then, using a blank of working buffer, lysozyme, ONPG, and Na_2_CO_3_, the reaction absorbances at 420 nm and 550 nm were determined. This process was repeated twice for each strain, using a new overnight culture inoculated with a unique colony, for a total of three biological replicates.

Miller Units were determined using the equation

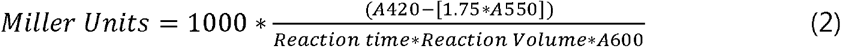

where 1.75 is an arbitrary value for correction for light scattering. However, we observed that the traditional equation often yielded negative values for controls, rather than the ideal of around 1 Miller Unit. We observed for our experiments, and in agreement with Krute *et al*. (50), the A_550_ correction coefficient of 1.75 was too large. After careful consideration, we chose to calculate Modified Miller Units using equation (2) but eliminating the A_550_ correction factor. This modified Miller unit value is what is reported as the average of three biological replicates, each taken as three technical replicates.

## RESULTS

### Crystallization of the B. subtilis azaaromatic (*yjdF*) riboswitch aptamer domain

To investigate the structural basis for ligand-dependent activation of the riboswitch, we determined crystal structures of the *Bsu yjdF* aptamer domain in complex with two representative ligands: chelerythrine, a known activator, and lumichrome, a non-activator. These functional assignments were established by Li et al, using a *lacZ* reporter assay in *B. subtilis* to evaluate riboswitch activity *in vivo* (17). We hypothesized that direct comparison of high-resolution structures of the aptamer domain in complex with an activating and non-activating ligand would reveal a mechanistic basis for differences in regulatory activity. To guide RNA optimization for crystallization trials, we analyzed the sequence conservation of 507 sequences from the current Rfam database (42). This analysis confirmed a pattern of nucleotide and secondary structural conservation consistent with prior findings (**Figure 1B**) (17). Additionally, we considered the in-line probing data of the *Bsu yjdF* RNA in the presence of chelerythrine (17) to avoid modifying RNA regions involved in ligand binding or vital tertiary interactions.

The conservation patterns indicate that multiple regions of the *B. subtilis yjdF* aptamer could be modified without impacting ligand binding activity. First, excluding the G•U wobble base-pair proximal to the central four-way junction, there is a lack of phylogenetic conservation in the L3 loop sequence and the P3 stem length, indicating that alteration of this element would not adversely affect ligand binding or gene regulation. Modest, unchanging reactivity by in-line probing at the L3 loop in the presence of several different ligands support this assessment (17). The 5’-side of the P4 helix and the L4 loop show a high sequence preference for pyrimidines, consistent with the hypothesis that the pyrimidine residues of L4 interact with the purine-rich RBS sequence to turn off gene expression (17). Arguing that L4 does not play a key role in establishing aptamer structure or ligand binding, the in-line probing of the isolated aptamer domain indicates that L4 displays modest reactivity and no change upon ligand binding. This analysis suggested that the lengths of P1, P3, and P4 and sequences of loops L3 and L4 (**Figure 1B**) can be varied without impacting ligand binding. In contrast, P2 and L2 display a very strong conservation pattern that is concentrated in L2 while nucleotides in this loop are protected from cleavage upon ligand binding (17). As such, alterations to the P2/L2 region were avoided due to its apparent functional importance. Given these observations, we engineered variants of the *Bsu yjdF* aptamer domain by replacing wild type L3 and L4 sequences with GAAA tetraloops, a common strategy in RNA crystallography (51), and by modifying the lengths of P1, P3, and P4.

To determine whether directed engineering of the *Bsu yjdF* aptamer for crystallography impacted ligand binding, we examined binding between mutants with those modifications and lumichrome. Lumichrome was chosen for its favorable properties relative to many other *yjdF* substrates. A fluorescence assay used previously served as the basis for our analysis of binding affinities (47). Titration of wild type aptamer RNA into a fixed concentration of lumichrome yields an observed equilibrium dissociation constant (K_D_) of 30±10 nM (**Figure 1C and 1D, Table 1**). Variants in which the wild type L3 or L4 loop is replaced by a GAAA tetraloop showed no change or very modest reduction of binding affinity (**Figure 1C and 1D, Table 1**). In contrast, replacement of the highly conserved wild type L2 sequence with a GAAA tetraloop reduces lumichrome affinity by >10,000-fold. These observations are consistent with our phylogenetic alignment and previous in-line probing data (17). To further validate these data, we assessed the binding of chelerythrine, an activating ligand, to the same set of RNAs and observed the same trend in binding affinities (**Table 1**). Thus, the P3/L3 and P4/L4 elements can be modified without impacting the ligand binding properties of the *yjdF* aptamer RNA.

**Table 1:** Binding of lumichrome to *B. subtilis yjdF*.

| **Mutation Group** | **RNA** | **K<sub>D</sub> (nM)<sup>a</sup>** | **K<sub>rel</sub><sup>b</sup>** |
| --- | --- | --- | --- |
|  | wild type | 30±10 | 1.0 |
|  | crystal RNA | 40±20 | 1.3 |
| **Terminal loops** | L2(GAAA) | >40,000 | >1290 |
|  | L3(GAAA) | 90±50 | 2.9 |
|  | L4(GAAA) | 30±20 | 1.0 |
| **P2a insertion** | +P2a | 40±10 | 1.3 |
|  | +P2a/U23A | 2,000±1,000 | 67 |
| **Pseudoknots** | U29C | 3,400±800 | 110 |
|  | A13G, U29C | 170±60 | 5.5 |
|  | G24C | 8,000±4,000 | 258 |
|  | G24C, C73C | 130±40 | 4.2 |
| **P4a** | U54A | 12±1 | 0.4 |
|  | C52G | 40±10 | 1.3 |
|  | U54A,C52G | 6±1 | 0.2 |
|  | G70C | 50±20 | 1.6 |
| **J1/2 - J3/4 interactions** | C12G | 1,200±200 | 39 |
|  | C12G, G48C | 800±400 | 26 |
|  | A11G | 360±30 | 12 |
|  | A11G, A11G, A50G | 1,400±100 | 45 |
| **Binding site triple** | A27G | >20,000 | >645 |
|  | A27G, U49C | >40,000 | >1,290 |
| **L2/backbone contact** | U23A | 1,800±800 | 58 |
|  | U23G | 9,000±1,000 | 290 |

<sup>a</sup>Buffer composition (Buffer A): 20 mM Tris-HCl, pH 7.5, 150 mM KCl, 5 mM MgCl<sub>2</sub>, 0.001% Nonidet P-40
<sup>b</sup>K<sub>rel</sub> = K<sub>D,mutant</sub>/K<sub>D,wild type</sub>

To find crystals of the *B. subtilis yjdF* riboswitch, we systematically varied the lengths of the P1, P3, and P4 helices and subjected each RNA variant to crystallization trials against a limited set of sparse matrices at 21 °C and 30 °C with a variety of ligands. One variant containing helix lengths of six, three, and five base pairs for the P1, P3, and P4 helices, respectively, yielded large crystals with lumichrome at 21°C over a period of 2-3 weeks and yielded diffraction to 2.55 Å. This RNA variant binds lumichrome with a nearly identical affinity to the wild-type RNA (“crystal RNA,” **Figures 1C** and **D, Table 1**). The same RNA was used to obtain crystals of a chelerythrine-*yjdF* complex. Final models of the lumichrome- and chelerythrine-bound *Bsu yjdF* aptamer domain included all 81 nucleotides of the RNA within the two protomers of the asymmetric unit (**Supplemental Table S2**). Below, we will focus our discussion of RNA structure on promoter A of the native lumichrome-*yjdF*, which is derived from the highest resolution dataset, and discuss the chelerythrine-bound structure where there are differences.

### The global architecture of the *yjdF* riboswitch is defined by two sets of coaxially stacked helices

The *Bsu yjdF* riboswitch adopts a T-shaped structure organized around two major axes. One axis of this fold includes the aromatic ligand binding pocket and is formed by P1, P2, and P3. The second axis projects from the central four-way junction into the distal regulatory helix P4/L4, which contains the anti-RBS sequence. These two helical axes are arranged nearly perpendicularly with the regulatory stack intersecting the ligand-binding stack near its midpoint. This interlocking architecture is stabilized by structured junctional regions and a network of non-canonical base-pairing interactions that scaffold the RNA into its folded conformation. Together, these features link the ligand-binding site to the regulatory region of the RNA, providing a structural framework for ligand-responsive gene regulation.

The four universally conserved helices of the *yjdF* family of riboswitches (**Figure 1B**, P1 – P4) are organized into two sets of coaxial stacks, comprising of P2-P3 and P3b-P4a-P4 (**Figure 2**, blue and orange, respectively) with the P1 helix collinear with the P2-P3 stack (pink, **Figure 2**). Notably, all of the base pairs predicted by the Breaker group were observed in the crystal structure (17). However, the crystal structure reveals unpredicted base-base interactions involving J1/2, J3/4, and J4/1 that form new structural elements: P3b, P4a, and two pseudoknots, PK1 and PK2 (**Supplemental Figure S4**). The orientation of the P1 helix is collinear to the P2-P3 stack, but its base pairs do not form a contiguous stack with the nucleotides in the P2-P3 stack. Instead, the first three nucleotides of J1/2 stack on P1 and anchor it to the P2-P3 via interactions through two A-minor triple interactions with two G-C pairs in the P2-P3 stack. This first coaxial stack contains the aromatic binding pocket and is discussed in further detail in the next sections.

**Figure 2.**
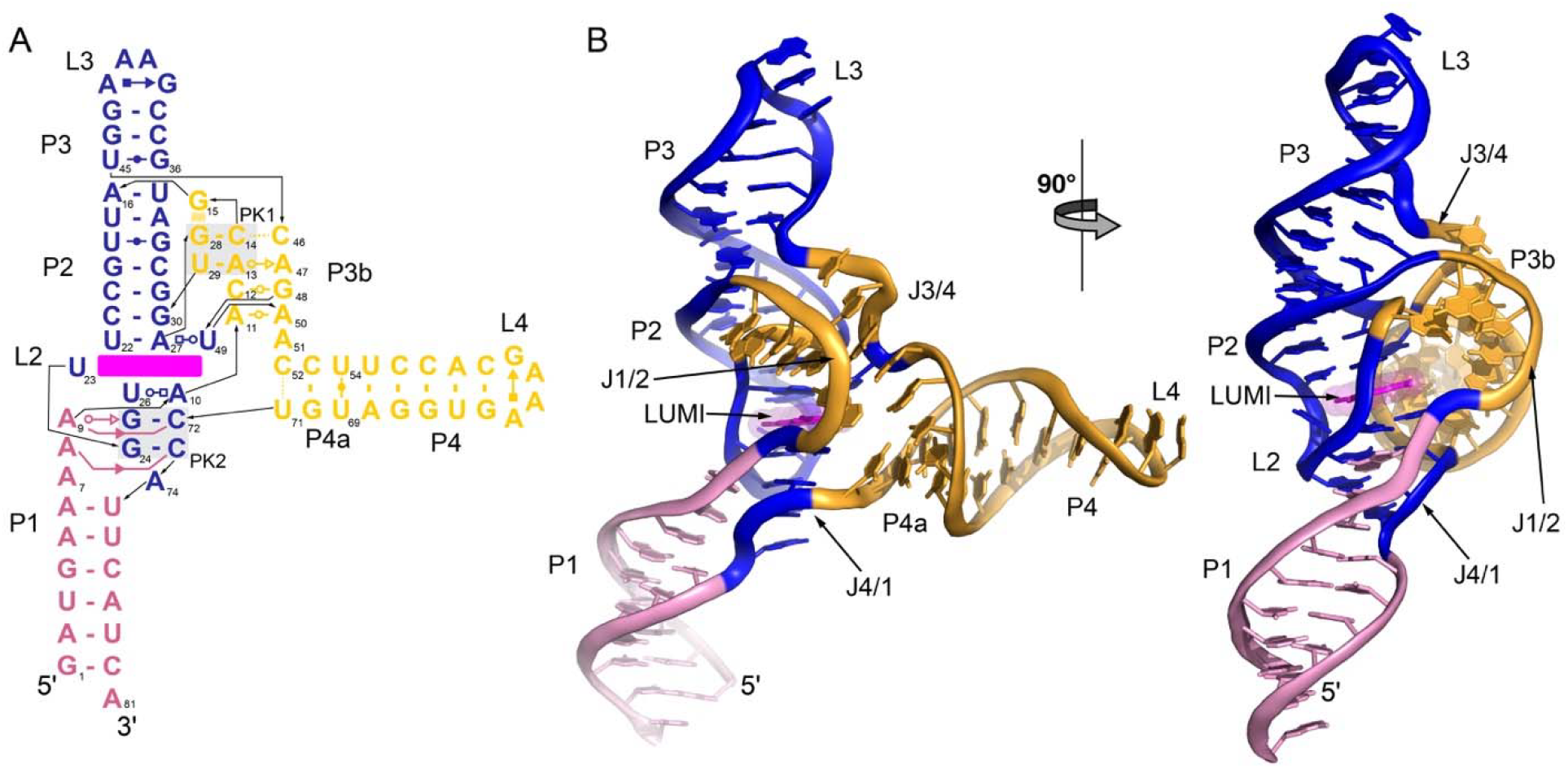
Architecture of the *B. subtilis yjdF* riboswitch aptamer domain. (A) Secondary structure of the RNA used for crystallization with base-mediated interactions denoted using the standard Leontis-Westhof notation (85). The numbering of the RNA is consistent with that used in the crystallographic models of the *B. subtilis yjdF* aptamer. The colors reflect the two coaxial stacks (blue and gold) and the auxiliary P1 helix (pink). (B) Cartoon representation of the three-dimensional structure of the *yjdF* aptamer using the same color scheme as in panel (A) and magenta for the ligand (lumichrome, LUMI). The left perspective highlights the three major joining regions (J1/2, J3/4 and J4/1) and the P3b-P4a-P4 coaxial stack (gold) while the 90° rotational perspective on the right highlights the P2-P3 coaxial stack (blue) and the relationship of the ligand (magenta) to the two coaxial stacks of helices.

The second coaxial stack spans nucleotides in J3/4 near the binding pocket to the terminal loop of P4 that contains the anti-RBS sequence. The structure suggests that this coaxial stack is functionally responsible for communication of ligand binding in the P2-P3 coaxial stack to L4 which contains the anti-RBS and thus mediates the regulatory response. The P3b-P4a-P4 coaxial stack is formed in significant part from base-base interactions between J1/2 and J3/4 (helix P3b) and a set of three base pairs on the junction-proximal side of P4 that were not predicted from phylogenetic analysis (helix P4a). The coaxial P2-P3 and P3b-P4a coaxial stacks are oriented at ∼145° angle with respect to one another with the P3b-P4a-P4 helix having a ∼50° bend centered around the intersection of P4a and P4 (**Supplemental Figure S5**). This kink in P4a may be induced by lattice contacts as the two protomers in the asymmetric unit interact with neighboring molecules through the engineered GAAA tetraloop of L4 (**Supplemental Figure S6**). This overall organization results in a general “T” shape architecture to the RNA as defined by the near-perpendicular arrangement (90° interhelical angle) of P4 relative to the P2-P3 coaxial stack (**Supplemental Figure S5**).

The global architecture of the *Bsu yjdF* riboswitch crystal structure is supported by phylogenetic, biochemical, and structural data. Phylogenetically conserved nucleotides are clustered around the ligand binding pocket and the P4a helix, which represent the functional centers of the aptamer domain (**Supplemental Figure S7A**). Similarly, the pattern of chelerythrine-dependent changes in in-line probing reactivity are localized near the ligand binding site and the P3b-P4a region (17) (**Supplemental Figure S7B**). Finally, the *Bsu* crystal structure closely aligns with the recently published crystal structure of the *Ruminococcus gauvreauii (Rga)* aptamer domain (52) and resembles the molecular envelope of the *Staphylococcus aureus* aptamer determined by small angle X-ray scattering (53). The *Bsu* and *Rga yjdF* aptamer crystal structures display near-identical overall global architecture with respect to the universally conserved P1, P2, P3, and P4 secondary structural elements. Superimposition of the backbone atoms of the conserved core of the *Bsu* and *Rga* chelerythrine-bound structures yield an RMSD of 1.37 Å (**Supplemental Figure S8**), indicating that these independently determined crystal structures of two variants of the *yjdF* aptamer domain are in general agreement.

The most notable difference between the *Bsu* and *Rga yjdF* structures is an additional variable helix (P2a) inserted into the 3’-side of J1/2 in the *Rga* variant. This additional helix, which is oriented parallel to P3, is present in 23% of the 507 sequences that we analyzed from Rfam. The presence of the P2a insertion is found predominantly in Clostridia and mostly deplete in Bacilli (**Supplemental Figure S9**). Using the *Bsu* and *Rga yjdF* structures as a guide, we constructed a *Bsu yjdF* variant in which appended the *Rga* P2a helix (**Supplemental Figure S10**). This variant binds lumichrome with a similar affinity to the wild type aptamer (30±10 versus 40±10 nM, respectively; **Table 1**), indicating that this insertion is likely neutral with respect to binding. While additional ligands remain to be tested, this result indicates that the P2a insertion alone is unlikely to define a functionally distinct subclass of the *yjdF* riboswitch as observed in the *ykkC* family (54,55).

### The P2-P3 coaxial stack hosts the aromatic binding pocket

The ligand binding site of the *yjdF* riboswitch is embedded within the P2-P3 coaxial stack, supported by the collinear P1 helix (**Figure 2b**). The P1 helix is formed exclusively by Watson-Crick-Franklin (WCF) base pairs with phylogenetic variations in length between five and seven base pairs. The first three nucleotides of J1/2 (A7, A8, and A9) cap the P1 helix and extend the stacking trajectory of the 5’-side of P1. Nucleotides A8 and A9 serve to anchor the P1 helix to the P2-P3 coaxial stack; A8 forms a type-II A-minor triple (56,57) with the G24-C73 WCF base pair, and A9 forms a type-I A minor triple with the G25-C72 WCF base pair (pairs *i* and *ii*, **Fig. 3A, B**). The phylogenetic conservation of these three adenosines is consistent with their observed structural roles. A7, which only engages in stacking interactions, is predominately a purine (91.3% A, 7.4% G, and <1% Y) whereas A8 and A9 are almost exclusive adenosine (98.8% and 99.4%, respectively), consistent with their roles in minor groove triple interactions. Interestingly, the equivalent of A8 and A9 in the wild type *Streptococcus ratti yjdF* riboswitch displays the strongest ligand-dependent 2-methylnicotinic acid imidazolide (NAI) chemical probing protection, suggestive of local conformational stabilization by the spatially adjacent azaaromatic ligand (53) (**Figure 2**).

**Figure 3.**
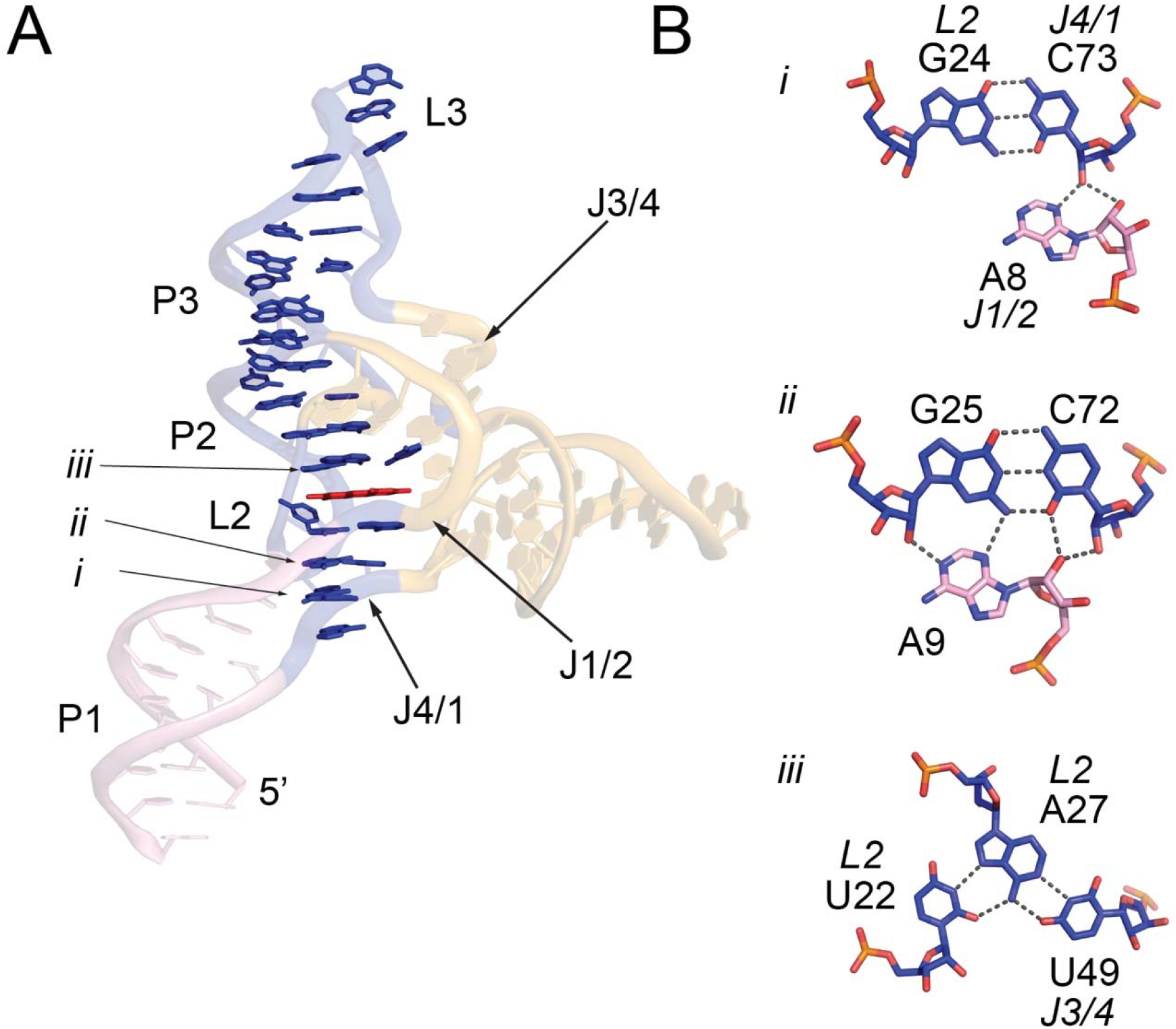
Structure of the “binding” P2-P3 coaxial stack. (A) Front view of the *yjdF* aptamer with the bases directly involved in stacking interactions between the P2 and P3 helices. Lumichrome is shown in red sticks in L2. Numbering *i*-*iii* denotes the positions of base-base interactions shown in panel (B). Panel (B) shows base-mediated hydrogen bonding interactions that form the PK2 element (*i* and *ii*) and part of the ligand binding pocket (*iii*).

The P2-P3 coaxial stack is formed by the predicted P2 and P3 helices stacking end-to-end with their terminal loops, L2 and L3, placed at each end of the stack (**Figure 2**). In this arrangement, L3 projects away from the core of the RNA and forms no contacts with the rest of the RNA. This is consistent with both high phylogenetic sequence variation in L3 and length of P3 as well as our observation that replacement of the wild type L3 with a GAAA tetraloop does not impact ligand binding. On the other side of the P2-P3 stack, L2 is anchored to the P1 helix via interactions with J4/1 and J1/2. Interactions between L2 and J4/1 form the two WCF G-C pairs of PK2 that form A-minor triples with A8 and A9 of J1/2 (see above). The identity of these two base pairs is almost invariant across phylogeny (98% and 98.4% for G24-C73 and G25-C72, respectively). Formation of PK2 is essential for high affinity azaaromatic ligand binding. A single point mutation in PK2 (G24C) significantly impairs lumichrome binding (K_rel_ = 260, **Table 1**) while the compensatory mutation (G24C, C73G) almost completely restores binding to near wild type affinity (K_rel_ = 4.2).

### Lumichrome and chelerythrine interact with a T-loop-like structural element

The azaaromatic ligand binding site is embedded primarily in L2 that participates in the P2-P3 coaxial stack (**Figure 3**). The structure of L2 fits within the U-turn consensus motif (58) that is a component of a broad array of loop motifs in biological RNAs including the T-loop (59,60). The consensus T-loop motif is defined by a sharp turn between nucleotides 2 and 4 that is part of the U-turn motif: a conserved purine at position 4 and a flanking non-canonical pair between nucleotides 1 and 5. The canonical T-loop motif generally has a space between nucleotides 4 and 5 for insertion of an aromatic group, such as the adenine moiety of a conserved adenosine in the tRNA D-loop (59). L2 in the *Bsu yjdF* riboswitch has a very similar structural arrangement (**Figure 4C, D**). Superimposition of the U-turn motif and the flanking U•A pair of tRNA^Asp^ (PDB ID 6UGG; (61)) and the *Bsu yjdF* L2 yields almost perfect structural homology (0.13 and 0.11 Å, respectively) (**Supplemental Figure S11**). The major difference between these two loops is that *yjdF* L2 has a single nucleotide insertion between canonical T-loop nucleotides 4 and 5 where the fourth nucleotide (U26) is in the same position as the D-loop adenine of the tRNA T-loop (**Supplemental Figure S11**). Lumichrome inserts itself between the expansion nucleotide (U26) and nucleotide 5 (A27) analogously to the tRNA D-loop/T-loop interaction. The backside of the binding pocket is created by U23 which makes a backbone contact to the non-bridging phosphate oxygen of U26, another hallmark of both the U-turn and T-loop motifs (58,59).

**Figure 4.**
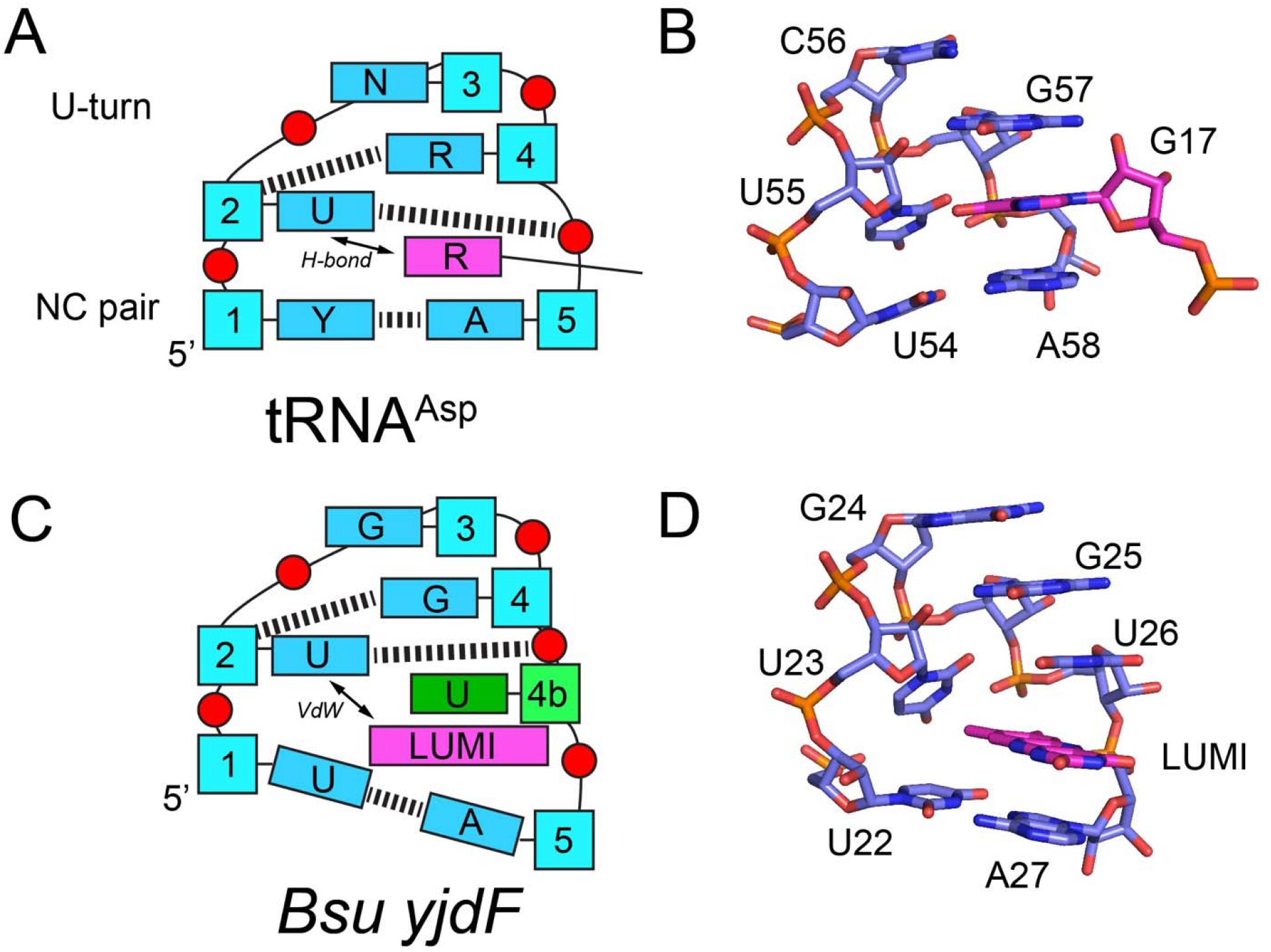
Structural homology between L2 of the *yjdF* riboswitch with the T-loop motif. (A) Schematic of a consensus T-loop from unmodified tRNA^Asp^ with R denoting purine, Y denoting pyrimidine, and N denoting any nucleotide. Dashed lines represent conserved hydrogen bonding interactions that serve to define the motif. Nucleobases in the T-loop motif are shown in blue, ribosyl moieties in cyan, phosphates in red, and an interacting adenosine nucleobase from the D-loop of tRNA shown in magenta. Nucleotides 2, 3, and 4 form a U-turn motif while positions 1 and 5 are involved in a non-consensus (NC) base pair. (B) Three-dimensional stick model of the T-loop of tRNA^Asp^; numbering is consistent with the crystal structure. (C) Schematic of L2 from the *Bsu yjdF* riboswitch bound to lumichrome (LUMI). Coloring and numbering are consistent with panel A except that the single nucleotide insertion is shown in green as position 4b. Bound lumichrome is represented as magenta. (D). Stick model of the T-loop of L2 of *Bsu yjdF* with bound lumichrome.

The expanded T-loop motif in L2 is embedded within a seven-nucleotide loop in which the two 3’-end nucleotides of the loop are bulged out to form long-range tertiary interactions. This is also a common feature of T-loop motifs found in the flavin mononucleotide (FMN) and cobalamin riboswitches where the two bulged nucleotides form interactions with a T-loop receptor motif (62). In the *Bsu yjdF* RNA, the last two nucleotides of L2 pair with nucleotides on the 3’-side of J1/2 to form the A13-U29 and C14-G28 pairs that comprise another pseudoknot (PK1). The C14-G28 pair of PK1 is nearly invariant (98% C-G, 0.8% U•G, and 1.2% other pairs) while the A13-U29 pair shows more covariation (83.8% A-U, 15.2% G-C, and <1% other pairs). A U29C mutation to destabilize PK1 is deleterious to lumichrome binding (K_rel_ = 110) while the compensatory mutation (A13G,U29C) restores binding (K_rel_ = 5.5), congruent with the importance of this pseudoknot in establishing RNA structure around the ligand binding site.

The azaaromatic binding site binds a single molecule of lumichrome, a non-activating compound, and chelerythrine, an activating compound in nearly equivalent positions within the T-loop motif of L2. The azaaromatic pocket is defined by the A10-U26 Hoogsteen/WCF base pair and the U22•A27-U49 base triple (pair **iii, Figure 3B**) and flanked on the backside of the binding pocket by U23 (**Figure 5**). The A10-U26 pair is nearly universally conserved in *yjdF* phylogeny (99.2% A-U and 0.8% other pairs) while the U22•A27-U49 is more variable (94.3% U•A-U, 4.5% C•A-U, 0.6% U•A-C, and less than 1% other triples). The overall architecture of this region of L2 creates a docking site for a planar azaaromatic compound, similar to the T-loop in tRNA that interacts with an adenosine in the D-loop (63) or the T-loop of the pyrimidine binding pocket of the thiamine pyrophosphate (TPP) riboswitch (64-66). Finally, it should be noted that in-line probing, dimethyl sulfate (DMS), and NAI (“SHAPE”) probing all suggest that this region is predominantly pre-organized in the apo-state (17,53), allowing for docking of azaaromatic compounds.

**Figure 5.**
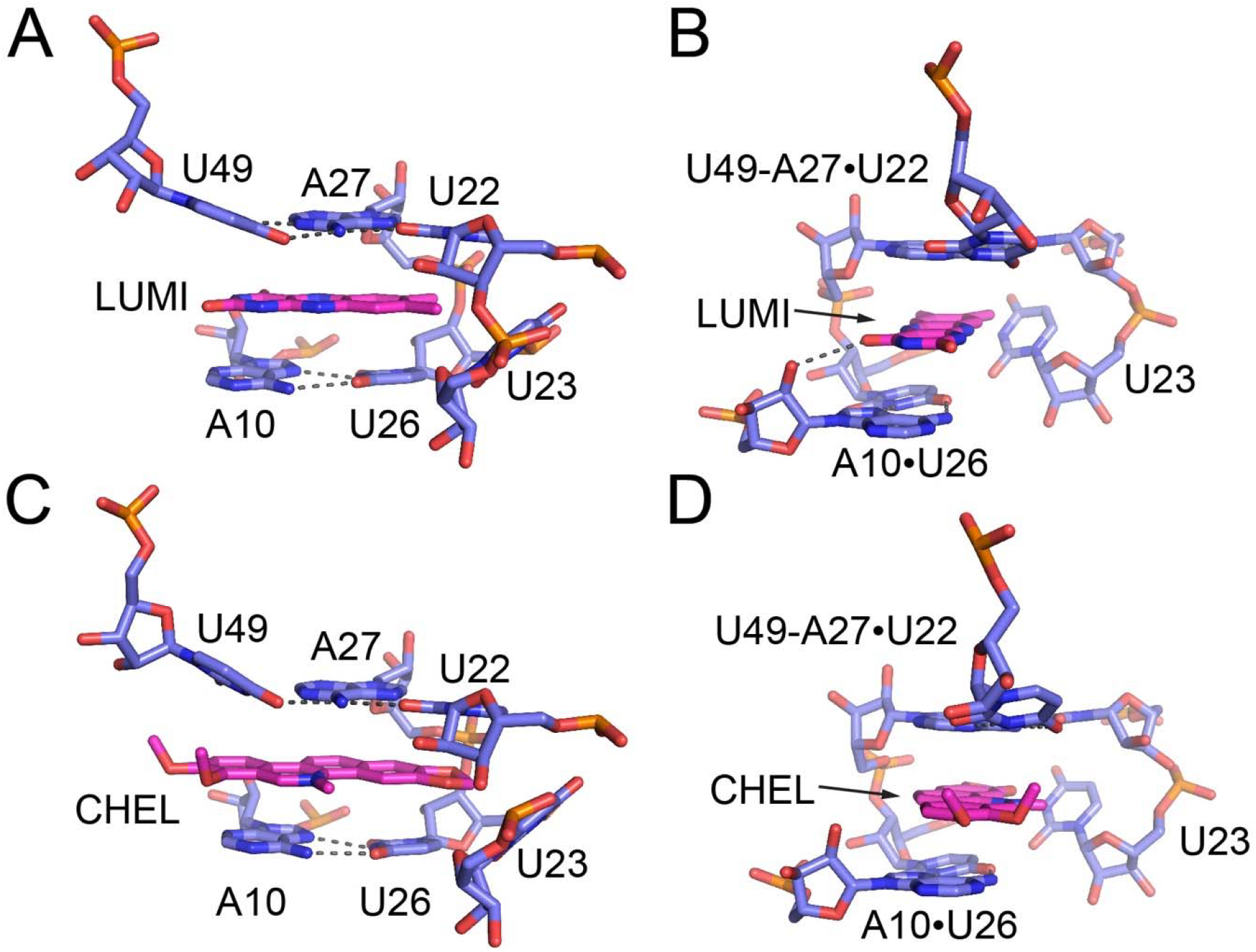
Structural details of ligand recognition by the *Bsu yjdF* riboswitch. (A) Front view of the binding pocket (blue) with bound lumichrome (LUMI, magenta). Dashed lines denote base-base hydrogen bonds and a LUMI-RNA hydrogen bond. (B) Side view (90° rotation of view in panel (A)) of the lumichrome binding pocket. (C) Front view of the ligand binding pocket (blue) bound to chelerythrine (CHEL, magenta). (D) Side view (90° rotation of view in panel (A)) of the chelerythrine binding pocket.

Part of the core of the T-loop motif is nucleotide position 2, which is important for stabilizing the U-turn of the backbone. The identity of this position in *yjdF* (U23 in *Bsu*) is variable (59.6% U, 26.6% A, 12.6% C, and 1.2% G), which distinguishes L2 from other T-loops that show a very strong preference for either uridine at this position in tRNA or guanosine in riboswitches (59). Across 193 structurally characterized T-loop motifs across biological RNAs, there are no examples of either adenosine or cytidine at this position (59). In the *Bsu yjdF* structures, U23 is positioned such that N3 hydrogen bonds to a non-bridging phosphate oxygen of U26 to form the backside of the pocket. In contrast, the equivalent of nucleotide 23 in the *Rga yjdF* structure is an adenosine that is flipped-out towards solvent, such that the azaaromatic pocket is open (52). Mutation of *Bsu* U23 to adenosine or guanosine is deleterious to lumichrome binding (K_rel_ = 58 and 290, respectively), indicating that the T-loop configuration with nucleotide 23 as a uridine promotes higher affinity ligand interactions. From an analysis of base conservation across phylogenetic variants with and without P2a, we observed that variants such as the *Bsu yjdF* that lack P2a strongly correlate with a uridine at position 23 while variants containing P2a such as the *Rga yjdF* mostly have an adenosine at this position (p = 3.6 × 10^−68^ and an odds ratio of 126.7 using a 2×2 contingency table and Fisher’s exact test for whether sequences with the P2a insertion have an adenosine at position 23). This could suggest that the A23 sequence variation in the context of *Rga yjdF* bearing the P2a helical insertion is not deleterious to azaaromatic ligand binding. However, when we placed the U23A mutation in the context of the *Bsu yjdF* P2a insertion mutant, we observe a 67-fold reduction in binding (+P2a/U23A, **Table 1**), which is the same loss of affinity as the U23A mutant in wild type *Bsu yjdF*. This indicates that the A23 sequence variant observed in 23% of *yjdF* riboswitches may represent a group of riboswitches with intrinsically lower affinity for azaaromatic compounds.

The ligand-RNA interaction mediated by L2 is dominated by π-stacking. Lumichrome is centered between the A10-U26 pair and U22•A27-U49 triple and forms a single hydrogen bonding interaction between a carbonyl oxygen (O4) and the 2’-hydroxyl group of A10 (**Figure 5B**). Since carbonyls are not common to all binding molecules identified by the Breaker group, this hydrogen bonding interaction is likely not critical for interaction with the RNA. Chelerythrine occupies a near-identical position in the binding pocket and does not appear to form any hydrogen bonding interactions with the RNA (**Figures 5C, D, Supplemental Figure S12**). The significantly higher affinity of *Bsu yjdF* for chelerythrine over lumichrome (**Tables 1 and 2**) is likely due to the positive charge in chelerythrine that creates a π-cation interaction with bases A10 and A27. Notably, the azaaromatic binding pocket is both extensive in size and open to bulk solvent on one side (**Supplemental Figure S13**), explaining why the Breaker group observed diverse planar azaaromatic molecules of varied sizes binding with nanomolar affinities. These compounds range from small two-member ring systems such as harmine and harmane to large five-member ring systems that include non-aromatic components as exemplified by staurosporine. Thus, this structure supports prior findings that the *yjdF* riboswitch can bind a diverse set of planar, azaaromatic compounds with high affinity, through what is likely to be a pre-organized hydrophobic pocket (17).

**Table 2:**
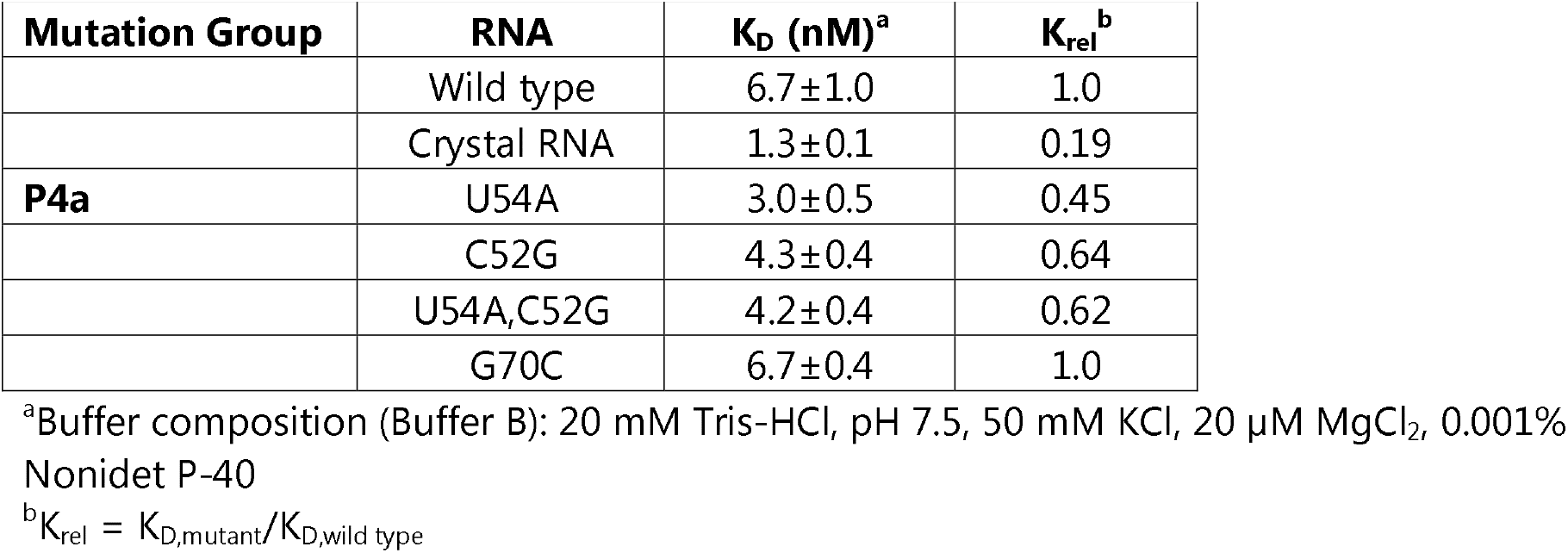
Binding of chelerythrine to *B. subtilis yjdF* in low salt conditions.

| **Mutation Group** | **RNA** | **K<sub>D</sub> (nM)<sup>a</sup>** | **K<sub>rel</sub><sup>b</sup>** |
| --- | --- | --- | --- |
|  | Wild type | 6.7±1.0 | 1.0 |
|  | Crystal RNA | 1.3±0.1 | 0.19 |
| **P4a** | U54A | 3.0±0.5 | 0.45 |
|  | C52G | 4.3±0.4 | 0.64 |
|  | U54A,C52G | 4.2±0.4 | 0.62 |
|  | G70C | 6.7±0.4 | 1.0 |

<sup>a</sup>Buffer composition (Buffer B): 20 mM Tris-HCl, pH 7.5, 50 mM KCl, 20 μM MgCl<sub>2</sub>, 0.001% Nonidet P-40
<sup>b</sup>K<sub>rel</sub> = K<sub>D,mutant</sub>/K<sub>D,wild type</sub>

### The *yjdF* aptamer domain binds and increases fluorescence of fluorogenic dyes

The structure of the *yjdF* riboswitch aptamer domain reveals a large and open binding pocket that accommodates a large array of flat, planar compounds. To further explore this, we examined the binding properties of a set of related compounds—fluorophores that have been shown to bind fluorescent light-up aptamers for live cell imaging (67). Most of these dyes fluoresce through conformational rigidification that is induced upon interaction with RNA (68-70). Development of these RNA imaging tools is an active area of research with many aptamers displaying modest performance, particularly in mammalian cells (71). It has been proposed that natural aptamers derived from riboswitches may yield better performance in cells due to their ability to rapidly fold with high fidelity (72).

Using fluorescent binding assays, we found that the *yjdF* aptamer domain can bind a variety of fluorogenic compounds, exhibiting a broad range of affinities with varying degrees of fluorescence turn-on or quenching *in vitro* (**Table 3**). Most notably, dyes that carry a positive charge (thiazole orange, chelerythrine, proflavine, 4′,6-diamidino-2-phenylindole (DAPI), and malachite green) display substantially higher binding, a trend which is also observed with chelerythrine versus lumichrome binding affinities. While *yjdF* is able to bind and activate the fluorescence of dyes that are currently used for RNA imaging in live cells such as HBC620 (Peppers aptamer (73)) and DFHO (Corn aptamer (74)), the most novel fluorophore binding is biliverdin. This compound, which is a product of heme degradation, has been used in fluorescence imaging of proteins that are tagged with smURFP (75), but lacks a known RNA aptamer that bind this compound. However, while the smURFP tag emission with biliverdin is in the far-red, *yjdF* RNA with biliverdin displays a broad increase in emission at a much shorter wavelength, around 490 nm. Nonetheless, these data together suggest that the *yjdF* aptamer could be engineered as a new RNA imaging tool potentially orthogonal to others currently in use (67).

**Table 3:** Fluorogen dye binding to wild type *B. subtilis yjdF* aptamer domain.

| **Fluorogen** | **K<sub>D</sub> (nM)<sup>a</sup>** | **Observed Fold Turn-on<sup>b,c</sup>** |
| --- | --- | --- |
| Thiazole Orange | 0.24±0.06 | 1,500±90 |
| Chelerythrine | 0.43±0.02 | 85±6 |
| Proflavine | 1.4±0.3 | 3.5±0.1 |
| DAPI | 4.5±0.9 | 55±2 |
| Malachite Green | 26±9 | 24.1±0.7 |
| Thioflavin-T | 39±7 | 1,950±80 |
| Biliverdin | 640±130 | 11.7±0.2 |
| HBC620 | 1,100±200 | 160±10 |
| NMMP | 3,900±100 | 17±1 |
| DFHO | 4,500±400 | 21±1 |

<sup>a</sup>Buffer composition (Buffer C): 20 mM Tris-HCl, pH 7.5, 135 mM KCl, 15 mM NaCl, and 1 mM MgCl<sub>2</sub>
<sup>b</sup>fold turn-on defined as the measured fluorescence in Buffer C of 1 μM dye in the presence of 10 μM RNA divided by the fluorescence of 1 μM dye in the absence of RNA.
<sup>c</sup>Respective wavelengths and bandwidths for measurements given in Table S3.

### The P3b-P4a-P4 coaxial stack connects the ligand binding site to the anti-RBS sequence

The P3b-P4a-P4 coaxial stack is dominated by base-base interactions that were not predicted from covariation analysis and provides a structural linkage between the ligand binding site in L2 and the regulatory anti-RBS sequence primarily in L4. The P3b helix (**Figure 6A**) is most closely associated with the ligand binding site as it is in close proximity and forms direct interactions with some components of the P2-P3 coaxial stack. The P3b helix is capped on one side by G15, which is the site of a turn in J1/2. Consistent with its role as an unpaired capping nucleotide, this position is predominantly a purine throughout phylogeny (71.6% A, 25.6% G, 2.2% U, and 0.6% C). Adjacent to G15 is PK1, formed by A13-U29 and C14-G28 interactions as discussed above. These two base pairs are supported by two nucleotides in J3/4: C46 and A47. C46 forms a single hydrogen bond between its N4 and the O2’ of C14 (pair **i, Figure 6B**). This interaction is supported by phylogenetic conservation pattern at this position (72.4% C, 26.0% A, 1.3% U, and 0.2%G) despite the N6 of an adenosine capable of supporting an analogous interaction. A47 forms a type-II A-minor triple with the A13-U29 pair has a higher conservation at this position (95.5% A, 2.6% deletion, 0.4% G, 1.0% C, and 0.6% U) (pair **ii, Figure 6B**). Adjacent to this pair is a trans WCF/WCF pair between C12 in J1/2 and G48 in J3/4 (pair **iii, Figure 6B**). This pair is highly conserved in phylogeny (97.2% C-G) and does not appear to be able to support either a trans WCF/WCF U-A pair (0.2%) or a trans wobble U•G pair (0.4%). The Y-R orientation of this pair is structurally important; a C12G mutation drops binding affinity for lumichrome by ∼40-fold, but the compensatory C12G/G48C mutation does not significantly restore binding (K_rel_ = 26, **Table 1**). The last pairing interaction of P3b is a trans WCF/WCF pair between A11 and A50, with A50 adopting a *syn* configuration (pair **iv, Figure 6B**). This pair is conserved as an A•A in 95.1% of *yjdF* sequences with 3.6% as the closely isosteric U•A pair. Mutations in this pair are moderately deleterious with an A11G mutation showing a 12-fold reduction in affinity and a A11G,A50G double mutant with a 45-fold reduction in ligand binding (**Table 1**). At the end of the P3b helix is an unpaired adenosine (A51) that links to the P4a helix and is only modestly conserved (82.4% A, 11.1% G, and 6.1% U).

**Figure 6.**
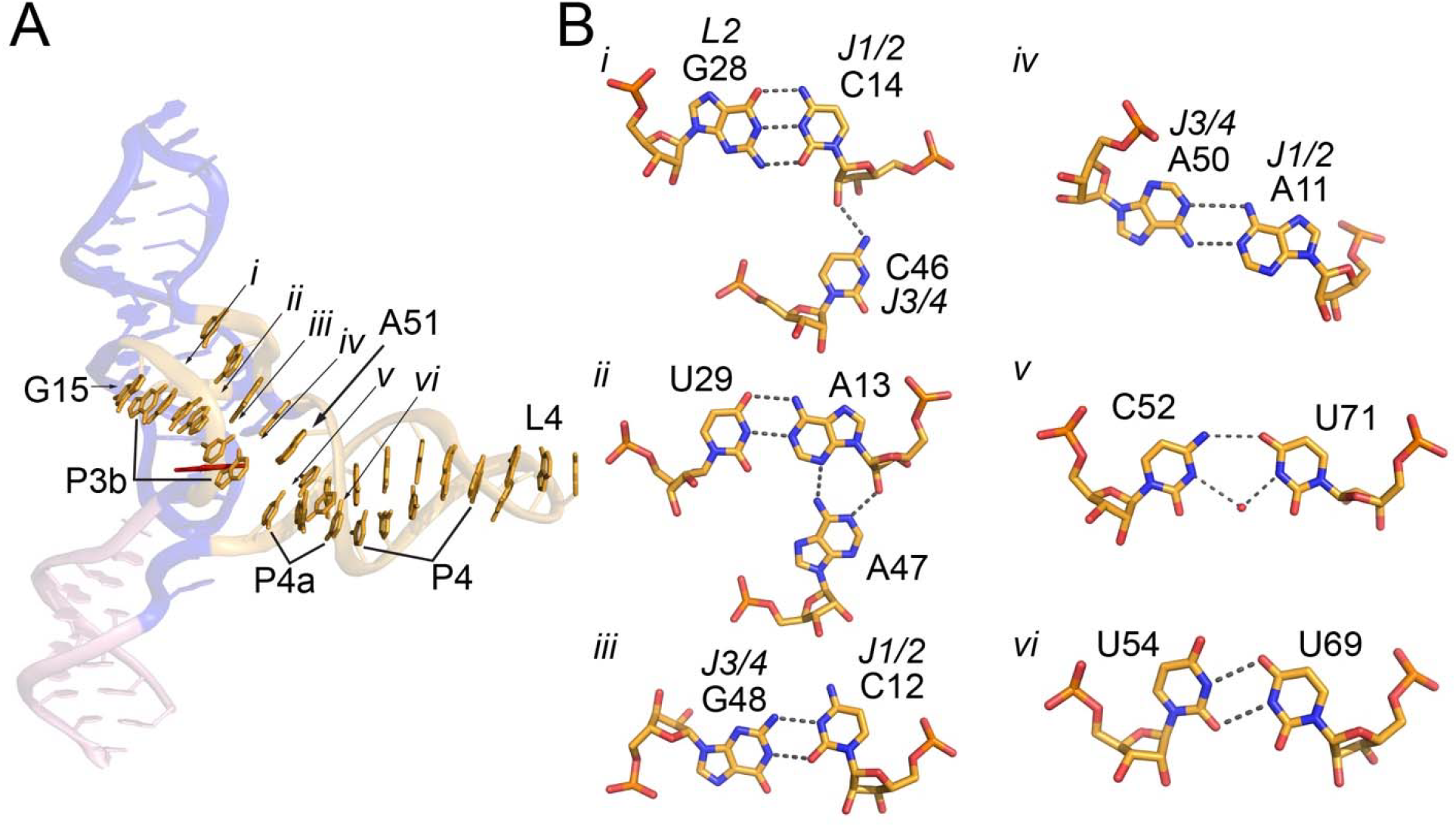
Structure of the “regulatory” P3b-P4a-P4 coaxial stack. (A) Front view of the *yjdF* aptamer with the bases directly involved in stacking interactions in the P3b-P4a-P4 coaxial helical stack (gold). Lumichrome is shown as red sticks. Numbering *i*-*vi* denotes the positions of base-base interactions shown in panel (B) along with highlighting unpaired nucleotides G15 at one end of the stack and A51 between the P3b and P4a elements. Panel (B) shows base-mediated hydrogen bonding interactions that form the P3b helix (*i* - *iv*) and the pyrimidine-pyrimidine pairs in P4a (*v* and *vi*).

Importantly, in the middle of contiguous stacking between C45 and A51 of P3b, a single nucleotide (U49) is extruded from the helix to form part of the triple that forms one side of the aromatic binding pocket (pair **iii, Figure 3B**), serving as a physical link between the ligand binding site and P3b. This is accomplished using the S-turn motif commonly found in many RNAs (76,77). The backbone of the S-turn motif is near the backbone of nucleotides U29 and C30 that are part of PK2 and P2, respectively (**Supplemental Figure S14**). In the structure, we observe two magnesium ions with one forming two inner-sphere contacts to non-bridging phosphate oxygens of C30 and U49 (**Supplemental Figure S15**). These ions are likely most responsible for the strong cation dependence of ligand binding. For example, the affinity of chelerythrine under high salt conditions is less than 10 pM (Buffer A) but drops ∼1000-fold when the cation concentration is dropped significantly (**Table 2**).

The second helical element of the P3b-P4a-P4 coaxial stack that was not predicted by phylogenetic analysis, P4a, contains two non-canonical pyrimidine-pyrimidine mismatches (C52•U71 and U54•U69) and a WCF C53-G70 pair. The first pair is a cis WCF/WCF interaction between C52 and U71 via a single hydrogen bonding interaction between the N4 of C52 and O4 of U71 (pair **v, Figure 6B**). This pair is further mediated by a well-ordered water interacting with the N3 of both C52 and U71. The high degree of phylogenetic conservation of this pair (97.4% C•U, 1% C•C, and 0.8% U•U) indicates that this pair is likely critical for riboswitch function. The second base pair of P4a is a WCF pair between C53 and G70 that displays a similar degree of high phylogenetic conservation (99.6% C-G, 0.2% C•U, and 0.2% C•A). The last pair is a wobble pair between U54 and U69. The O2 of U54 hydrogen bonds with N3 of U69 while the N3 of U54 hydrogen bonds with the O4 of U69 (pair **vi, Figure 6B**). In contrast with the other two pairs in the P4a helix, the conservation pattern at these positions is more variable but does require a pyrimidine-pyrimidine pairing (69.4% U•U, 15% U•C, 12.4% C•C, 1.4% C•U, and 1% G•U). This last pair is adjacent to the first base pair of P4 (U59-A68), forming a contiguous helical element. It should be noted that this helix is the center of a ∼55° bend in the P3b-P4a-P4 coaxial stack and may serve as a flexible hinge between the body of the aptamer containing the ligand binding site and the distal L4 regulatory loop. While this bend angle may be influenced by lattice contacts mediated by L4 (**Supplemental Figure S5**), a similar local bend in P4a is observed in the *Rga yjdF* crystal structure, despite also being potentially influenced by lattice contacts (52). Reinforcing the importance of P4a, many of the nucleotides show a chelerythrine-dependent protection in in-line probing experiments (**Supplemental Figure S6**) (17).

To assess the impact of P4a on ligand binding, we examined the impact of a set of mutants on both lumichrome and chelerythrine binding. The first set of mutations, C52G and U54A, were designed to stabilize the P4a helix by creating a G•U and A-U pair, respectively. This was complimented by the C52G,U54A double mutant that further stabilizes the P4a helix. The G70C mutation, on the other hand, was designed to further destabilize the P4a helix by creating a C•C mismatch. The stabilizing mutants have, at best, a modest impact on ligand binding (**Tables 1 and 2**). For example, the U54A mutation increases binding affinity to lumichrome by ∼2.5-fold while the double mutant (U54A,C52G) increases by ∼5-fold. Chelerythrine binding to these mutants showed similar but modest improvement in binding affinity. Destabilization of the P4a helix with the G70C mutation also had a very small or no impact on binding for either ligand. These data demonstrate that while there is a high degree of conservation of nucleotide and base pair identity in P4a, the identity of these base pairs has little impact on binding of these azaaromatic compounds to the L2 site.

### P4a is critical for regulatory activity but suggests yjdF may still be an orphan riboswitch

Structural analysis of the *Bsu* and *Rga yjdF* riboswitch aptamer domains revealed three unexpected base pairs that form P4a that extend the P4 helix. This observation suggests a hypothesis for regulation of gene expression by the *yjdF* riboswitch: ligand-dependent formation of P4a prevents the anti-RBS in L4 from interacting with the RBS to enable translation (**Figure 7**). To further explore this hypothesis, we created a *B. subtilis* reporter in which the *yjdF* riboswitch is immediately upstream of the *lacZ* gene under control of the IPTG-inducible Pspank promoter. This reporter in the presence of IPTG and X-gal shows little expression of the *lacZ* reporter gene, but in the presence of 50 nM chelerythrine, we observe an increase in β-galactosidase activity as measured using a Miller assay (**Figure 8A**). This chelerythrine-dependent activity is similar to what was observed previously (17). As a control, we also made a *yjdF* reporter where the wild type L4 sequence was replaced with a GAAA tetraloop to disrupt the anti-RBS/RBS. This mutant displays significant β-galactosidase activity in the absence of ligand (**Figure 8A**). Further, we tested the non-activating compound lumichrome which failed to induce activation at 5 µM and only weakly induces activation at 50 µM. These results are consistent with prior data and reinforce the distinction between chelerythrine and lumichrome as activating and non-activating compounds, respectively. (**Figure 8B**).

**Figure 7.**
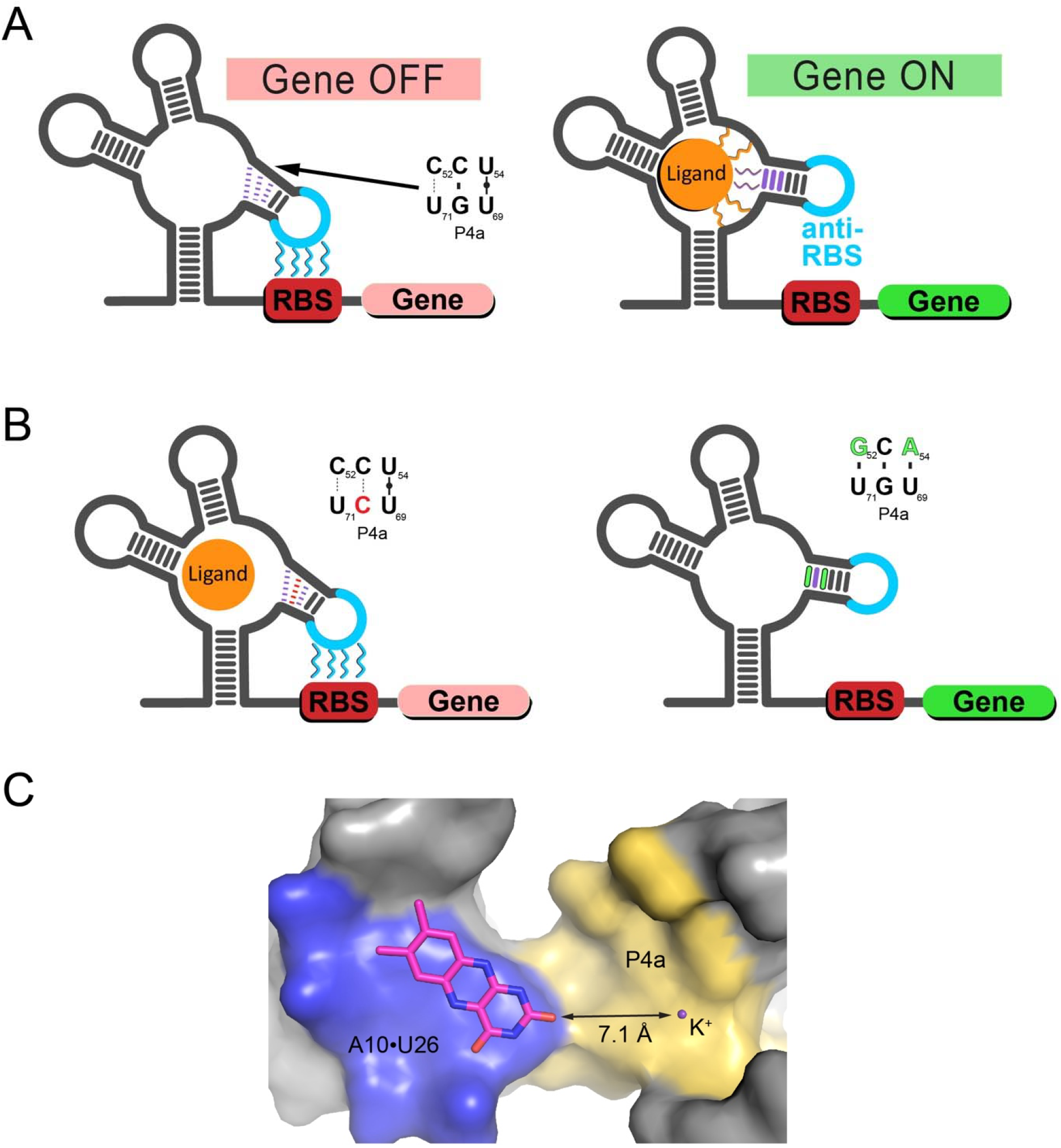
Cartoon of aspects of the regulatory mechanism of *yjdF* riboswitches. (A) The wild type *yjdF* riboswitch acts as a genetic ON switch. In the absence of ligand, the anti-RBS sequence in L4 (cyan) can pair with the RBS to block translation. In this state, it is proposed that the weak Y-Y pairs in P4a do not form. In the presence of an activating ligand, the pairs of P4a form and rigidify the P4 helix in an orientation that prevents it from pairing with the RBS and thereby promoting translation. (B) The effect of ligand binding is proposed to be mimicked by destabilizing (left) or stabilizing (right) mutations in P4a. (C) Surface representation of the aromatic binding pocket (blue) emphasizing its continuity and proximity to the major groove side of the P4a helix (yellow). K^+^is a modeled potassium ion sitting in the major groove between the C52-U71 and C53-G70 pairs.

**Figure 8.**
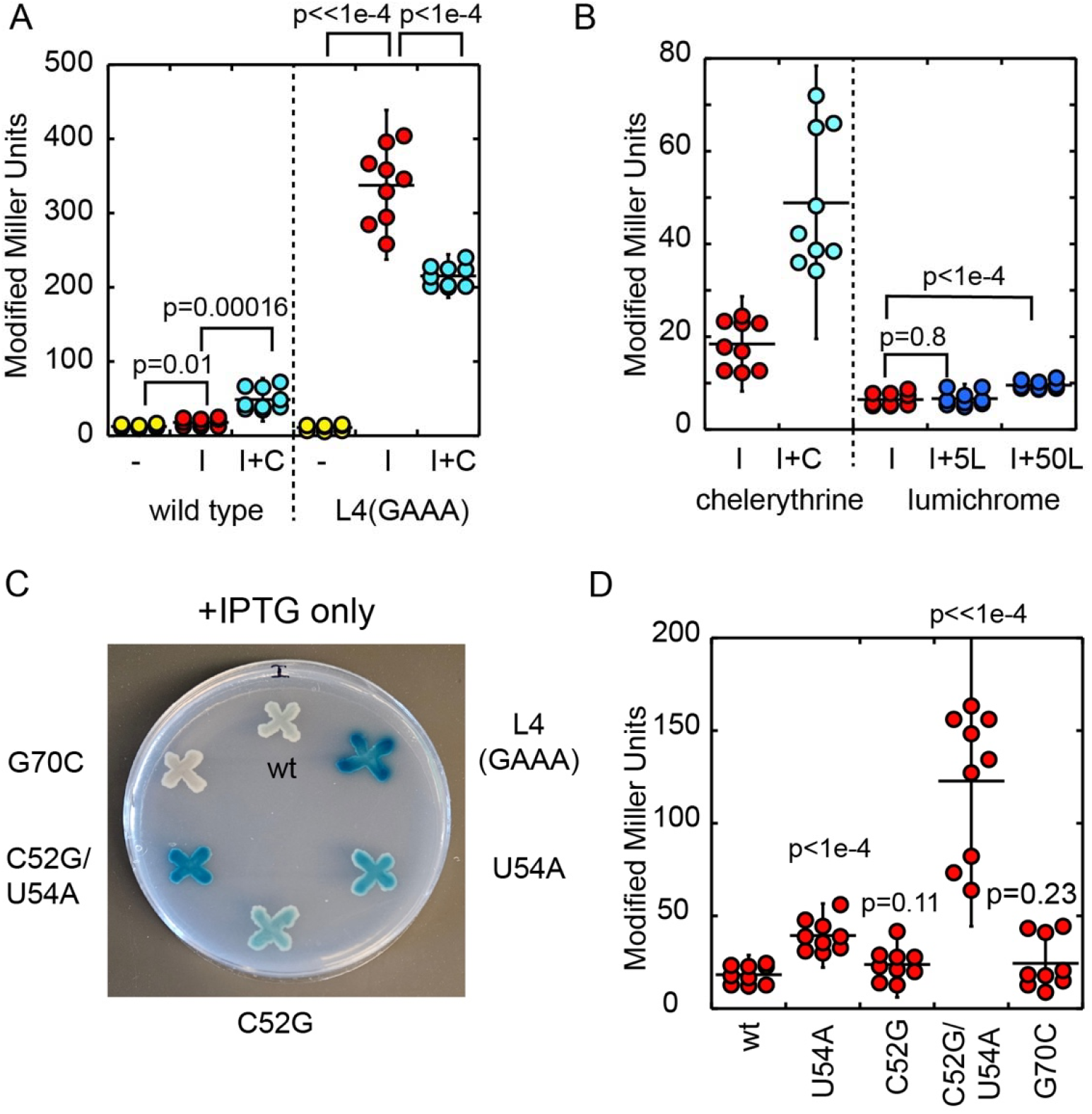
P4a stabilizing or destabilizing mutations affect expression in the absence of an activator ligand. (A) The wild type *yjdF* riboswitch can induce beta-galactosidase activity when *B. subtilis* is grown in the presence of 50 nM chelerythrine. The *yjdF*-lacZ transcript is under IPTG (I) control via the Pspank promoter. In the absence (yellow) and presence (red) of 1 mM IPTG, low beta-galactosidase activity is observed using a Miller assay. Addition of 50 nM chelerythrine with 1 mM ITPG (I+C, cyan) yields a modest increase in reporter activity. A mutation in L4 that converts it to a GAAA tetraloop that cannot pair with the RBS shows strong induction of LacZ expression upon addition of IPTG alone and a small, systematic decrease in that expression upon addition of chelerythrine. (B) The first two sets of data are the same as the wild type (I) and (I+C) in panel (A), while the right side shows reporter activity in the presence of IPTG and either 5 µM (I+5L) or 50 µM (I+50L) lumichrome. At 50 µM lumichrome, we observe a distinct but modest activation in comparison to 50 nM chelerythrine. (C) Defined-medium/agar plate of various *B. subtilis yjdF-LacZ* reporters in the presence of IPTG alone for 48 hours. P4a mutations in the *yjdF* aptamer are denoted to the side that either stabilize (U54A, C52G and C52G/U54A) or destabilize (G70C) the helix. (D) Miller assay of a set of the same set of mutants shown in panel (C) to quantify the degree which mutations in P4a repress or activate LacZ expression. In this figure, error bars correspond to two standard deviations from the mean and the p-values were calculated using a student’s two-tailed t-test using Excel.

To examine the role of P4a in activation of gene expression, we examined a set of mutants that are expected to either stabilize or destabilize this element (**Figure 7B**). These mutants were shown to have a very modest effect on both lumichrome and chelerythrine binding to the aptamer domain (**Tables 1 and 2**). Single point mutations expected to stabilize the P4a helix (U54A and C52G) show enhanced *lacZ* activity in the absence of ligand as compared to wild type (**Figure 8C, D**). Combining these two mutations to form three WCF pairs in P4a yield a strong enhancement of *lacZ* activity in the absence of ligand. Conversely, destabilization of the G-C pair in P4a by conversion to a C-C mismatch (G70C) yields no *lacZ* activity, like wild type. Thus, we can mimic ligand-dependent activation of gene expression by altering pairing in the P4a helix, supporting the hypothesis that stabilization of P4a makes the anti-RBS sequence in L4 inaccessible to the RBS, thereby activating translation. This panel of mutants were also tested for their ability to be activated in the presence of 50 nM chelerythrine (**Supplemental Figure S16**). Both U54A and C52G displayed strong ligand-induced gene expression with a degree of activation similar to wild type. Conversely, the G70C mutation that destabilizes P4a showed no chelerythrine-dependent activity.

Crystal structures of the *yjdF* riboswitch bound to chelerythrine suggest that ligand-dependent stabilization of P4a occurs indirectly via conformational changes in the P3b that get communicated through the P3-P4a-P4 coaxial stack. If the ligand indirectly acts upon the P4a helix, chelerythrine-dependent activation should be strongly affected by changes in nucleotide and base pair identity. However, the above data suggests that at the C52G and U54A mutations do not drastically affect chelerythrine-dependent activation. To further probe the influence of sequence in P4a on regulation, we sought to test mutations that would not severely affect P4a stability. Examination of the phylogenetic alignment of 507 *yjdF* riboswitch sequences, it is observed that the C52•U71 and C53-G70 base pairs are conserved >98% and that transversions of these two pairs are rarely, if ever, observed. This indicates that the polarity of these two pairs is important for riboswitch function and seemingly inconsistent with an indirect communication model, which would likely not be sensitive to the base pair orientation. Examination of the two transversion mutations (C52U,U71C and C53G,G70C) shows that the two transversions still show activation of translation by chelerythrine (**Figure 9**) with the C53G,G70C mutation displaying activity like wild type. This suggests that if azaaromatic compounds are the biologically relevant activators of the *yjdF* riboswitch, covariation patterns of the C-U and C-G pairs should be observed. A further mutation (C53U,G70A) that creates a weaker A-U pair shows no ability to activate lacZ expression, consistent with the G70C mutation.

**Figure 9.**
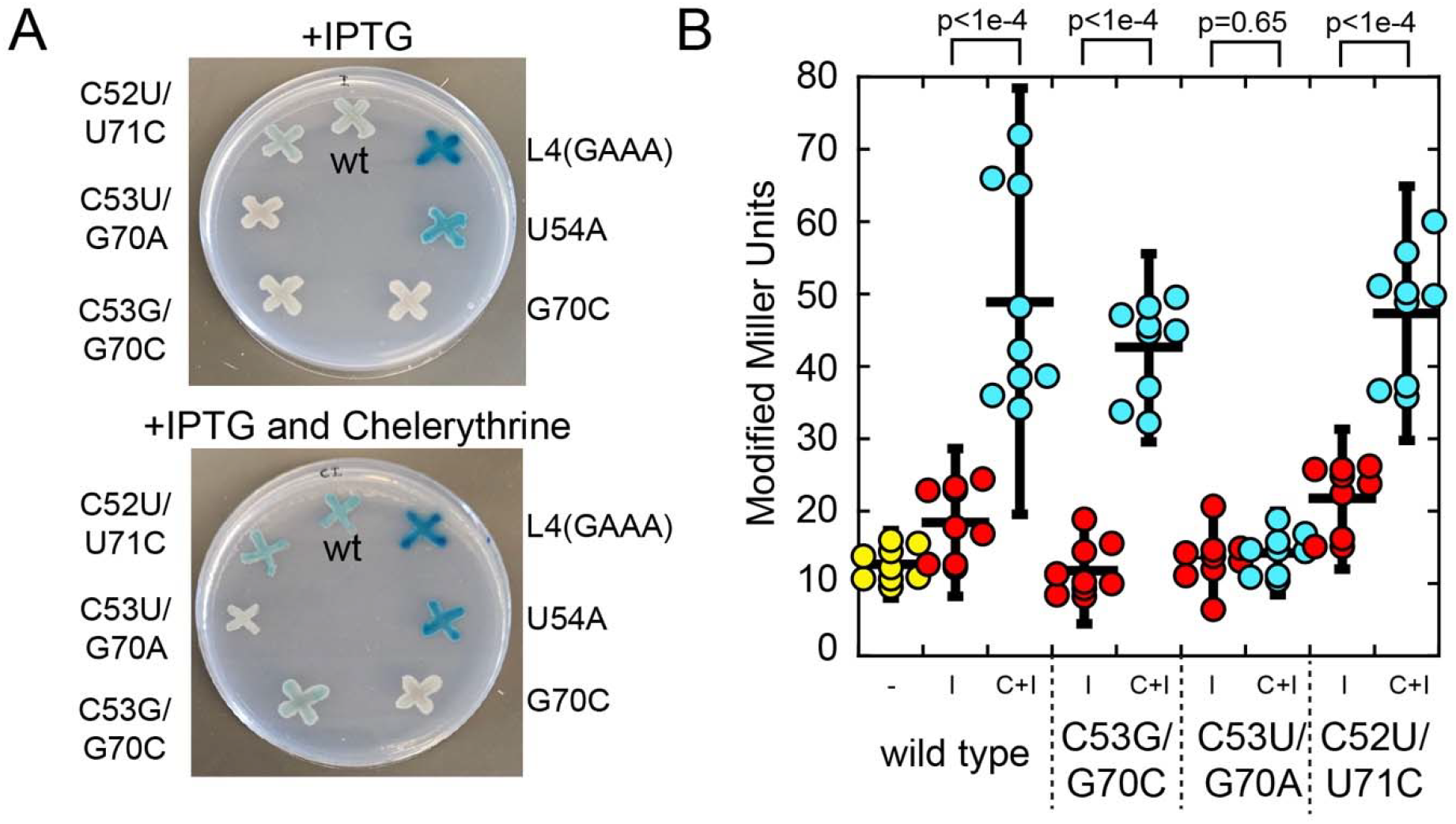
Transversions of highly conserved base pairs in P4a do not affect chelerythrine-dependent regulation by *yjdF*. (A) Define-medium/agar plate of various *B. subtilis yjdF-LacZ* reporters in the presence of IPTG alone (top) or IPTG plus 50 nM chelerythrine (bottom) for 48 hours. Identity of *yjdF* variant for each reporter given to the side. U54A and G70C mutations are plated for direct comparison to the pair mutations. (B) Miller assay quantifying the effect of base pair mutations on chelerythrine dependent regulation. The first three data (wild) type are the same as in panel (A) of Figure 8. Mutations that transverse the C53-G70C or the C52U/U71C show wild type chelerythrine dependent activation (red versus cyan) while a C53-G70 to U53-A70 base substitution is strongly deleterious, like the G70C mutation. In this figure, error bars correspond to two standard deviations from the mean and the p-values were calculated using a student’s two-tailed t-test using Excel.

These data suggest a “direct contact” hypothesis in which the cognate ligand interacts with functional groups in the major groove of P4a via hydrogen bonding interactions, in a mode of RNA-ligand recognition that would be highly sensitive to the spatial arrangement of the bases. Examination of the lumichrome-*yjdF* structure reveals a potassium ion sits in the major groove between the C52-U71 and C53-G70 base pairs is only ∼7 Å away from a carbonyl oxygen of lumichrome (**Figure 7C**). This suggests that a metabolite in the binding pocket could have a functional group(s) that extends to the major groove of P4a and directly contacts the highly conserved nucleotides to directly manipulate the conformation of P4a and, thereby, the anti-RBS/RBS interaction.

## DISCUSSION

The *yjdF* aptamer domain is unique amongst the characterized riboswitches for both its ligand binding promiscuity and unclear criteria of the chemical features that enable translational regulation. To understand these outstanding questions, we performed a detailed structural and functional analysis of the *B. subtilis yjdF* riboswitch that complements prior work from the Breaker group (17). Our structures of a non-activating (lumichrome) and activating (chelerythrine) compound in complex with the *Bsu yjdF* aptamer domain reveal a large binding pocket that accommodates a broad spectrum of planar, aromatic compounds, consistent with structures of the *R. gauvreauii yjdF* RNA bound to two different activating compounds (52). Our analysis emphasizes that both activating and non-activating compounds occupy the same site in the RNA providing no clear explanation for why certain compounds bind and regulate while closely related compounds bind tightly but do not regulate. Functional data from a *lacZ* reporter in *B. subtilis* reinforce that, while chelerythrine can regulate, mutations to a highly conserved element that is crucial for regulation (P4a) do not affect chelerythrine-dependent translational activation. These data highlight our lack of understanding of the role of the almost universally conserved base pairs in the P4a helix in the function of the *yjdF* riboswitch.

Structural analysis of the *Bsu* and *Rga yjdF* riboswitches provides a clear understanding of its degenerate specificity of ligand binding. The *yjdF* riboswitch creates a large binding pocket primarily situated in L2 whose shape supports binding of diverse-sized planar compounds with a preference for aromatic systems containing a positive charge. This binding pocket is likely mostly preorganized for ligand recognition. In-line probing data shows weak ligand-dependent protections in nucleotides surrounding the pocket with stronger protections around P3b (17). SHAPE and DMS chemical probing reveals even fewer protections in the *S. rattus yjdF* riboswitch with the only ligand-dependent changes observed around the equivalent position of A8 and A9 (53). The aromatic binding pocket, flanked by a WCF/H U•A pair and a U•A-U base triple, is similar to the Peppers aptamer and the xanthene binding site of the FMN aptamer (**Supplemental Figure S17**) (78-80). The Peppers aptamer binds a series of HBC fluorophores, none of which bear a common single function group but rather are related by being aromatic systems (73). The similar promiscuous binding behavior of the Peppers aptamer allows it to bind a variety of fluorophores with different excitation and emission wavelengths, enabling the same aptamer to be used to image a spectrum of colors. Similarly, the *yjdF* riboswitch can also bind an array of fluorophores, some of which are promising leads for the development of this RNA as a new imaging tag. Overall, our structure and associated binding data clearly explain the degenerate specificity of the *yjdF* riboswitch for compounds bearing aromatic systems.

While we expected the crystal structures of the *yjdF* aptamer in complex with an activating and non-activating ligand to reveal different binding modes, our data cannot reconcile the ability of only a small subset of binding compounds to activate translation (48). Both the lumichrome and chelerythrine structures are nearly identical at both a global and local structure with an RMSD of 0.58 Å and 0.50 Å over 80 nucleotides for protomers A and M of each, respectively. However, it is clear from our *yjdF*-regulated *lacZ* reporter assay in *B. subtilis* that chelerythrine induces translation, consistent with data from the Breaker group (17). Structures of the *Bsu* and *Rga yjdF* riboswitches in complex with chelerythrine indicate that communication between the ligand occupied L2 and regulatory L4 would likely be the result of structural changes in P3b and P4a upon ligand binding. In this mechanism, the P4a element acts as a “transducer element” that serves as an intermediary between the ligand binding pocket and the anti-RBS in L4 (52). In-line probing of the *Bsu yjdF* riboswitch aptamer domain in the absence and presence of chelerythrine revealed moderate ligand-dependent protections throughout PK1, P3b, and P4a (**Supplemental Figure S7B**) (17). This suggests that in the absence of an activating ligand the region around P3b/P4a is dynamic, enabling L4 to interact with the RBS to create a base pairing interaction that would occlude the ribosome from translating the message. In the presence of ligand, the P3b-P4a-P4 coaxial stack becomes more rigid, preventing L4 from accessing the RBS, and thereby enabling translation. The ability of stabilizing mutations in the P4a helix to promote translation in the absence of ligand supports a mechanism in which localized conformational changes in P4a drive regulatory activity. This proposed mechanism is like that of the preQ_1_-III riboswitch in which ligand binding to the central core is transmitted to a peripheral loop whose anti-RBS interacts with the RBS to regulate translation (81).

Our structural and functional analyses lead us to propose that the biologically relevant cognate ligand of the *yjdF* riboswitch remains unknown. There are a few critical observations that suggest an alternative effector molecule exists. First, the structures of the non-activating lumichrome and activating chelerythrine are almost identical, providing no clear insights into how an azaaromatic compounds have a differential effect upon the L4-RBS regulatory interaction. We also note that the non-activating compound riboflavin, which shares the same xanthene ring system as lumichrome, induces a similar set of protections by in-line probing, albeit at higher ligand concentrations. This indicates that ligand-induced conformational changes in the RNA for activators and non-activators is similar (17). Second, comparison of the position of chelerythrine in the *Bsu* and *Rga* riboswitch structures indicates that orientation of the ligand is not specific—that is, the positioning of the dioxolane ring can be either towards nucleotide 23 of L2 or away from it (48). Sanguinarine, which does not activate, differs from chelerythrine only by the presence of two dioxolane rings at each end. If the methoxy groups of chelerythrine that replace one of the dioxolane rings of sanguinarine are crucial for their difference in regulatory activity, it remains unclear how the promiscuity of the chelerythrine configuration in the binding pocket promotes activation. Third, transversion mutations in P4a that are phylogenetically forbidden do not appear to impact regulation by chelerythrine. If these pairs strongly affect regulation, as suggested by the stabilization and destabilization mutants, we expect the transversion mutants to significantly abolish chelerythrine-dependent regulation. Fourth, while the *in vivo* reporter data demonstrate that *yjdF* induces translation when cells are grown in the presence of chelerythrine and other select compounds, a direct association between the ligand binding event and gene activation has not been established. There remain potentially confounding variables surrounding ligand permeability, transport, and metabolism. Many of these azaaromatic compounds are also toxic to bacteria, and they could induce a stress response that triggers formation of another common metabolite that is the true effector molecule. Addition of 50 nM chelerythrine to the constitutively active L4(GAAA) mutant always results in lower Miller unit activity relative to the no ligand control, despite controlling for cell density (**Figure 8B**). This is likely the result of chelerythrine’s known capacity to hinder bacterial protein synthesis and disrupt the cell wall (82). Finally, we note that a destabilizing variant at position 23 in the binding pocket (A23) is highly phylogenetically represented. This suggests that molecular interactions with the aromatic pocket may not be the sole driver of the full set of interactions of the natural ligand that would drive both high affinity and activation of translation. Taken together, these structural and functional analyses of the *yjdF* riboswitch with activating and non-activating ligands fail to provide a compelling mechanism for regulation by a diverse group of azaaromatic compounds.

Instead, we propose that the biological effectors of this RNA are not purely azaaromatic compounds but rather some other endogenous metabolite. Biochemical and structural analysis of this RNA unambiguously demonstrate that the cognate ligand must contain a planar system as indicated by the large planar pocket formed by a set of universally conserved nucleotides in L2 and the joining regions. However, the major groove face of P4a is situated ∼7-8 Å away from lumichrome in our structure (**Figure 7C**), suggesting that a metabolite containing additional non-aromatic chemical groups could extend into this region to directly stabilize the P4a helix and promote translation of the mRNA. Such bipartite recognition in riboswitches is common. For example, the FMN riboswitch contains a flat, planar pocket for the aromatic xanthene ring and a phosphate binding pocket formed by a set of highly conserved unpaired guanosines (83). The ribosyl sugar that links the xanthene ring and phosphate group is not recognized by the RNA and can be substituted with other flexible linkers (83,84). We propose the cognate ligand of this RNA is like FMN in that it is composed of both aromatic and non-aromatic moieties directly contacted by the RNA to achieve the appropriate regulatory response. In this direct mechanism, the ligand would contact the highly conserved C52,C53/G70,U71 nucleotides to prevent pairing between the anti-RBS of L4 and the RBS. This model does not preclude a different base pairing arrangement or helical structure of P4a from that observed in either the current *Bsu* or *Rga yjdF* crystal structures, as they are likely influenced by lattice contacts. This hypothesis suggests the highly conserved P4a element is part of the effector binding pocket and not a transducer element. Thus, we believe that our work suggests that the *yjdF* riboswitch remains an orphan whose biologically relevant effector remains to be discovered. This would also indicate that the *yjdF* riboswitch, rather than being a promiscuous or degenerately specific regulatory element that responds to a spectrum of chemically related compounds is responsive to a single metabolite or several closely related metabolites, like every other known riboswitch.

## Supporting information

All Supplemental Data

## Data Availability

Atomic coordinates and structure factor amplitudes have been deposited into the Protein Data Bank (PDB) database under accession codes 9EC4, 9EBP, and 9EBV.

## Supplementary data

Supplementary data is available at NAR online.

## Acknowledgements

We thank Dr Jay Nix and the staff of beamline 8.2.2. of the Advanced Light Source, Lawrence Berkeley National Laboratory, for their support with remote crystallographic data collection. Beamline 8.2.2. of the Advanced Light Source, a DOE Office of Science User Facility under Contract No. DE-AC02-05CH11231, is supported in part by the ALS-ENABLE program funded by the National Institutes of Health, National Institute of General Medical Sciences, grant P30 GM124169-01. We thank the Macromolecular X-ray Crystallography Core (RRID:SCR _ 019310) at the University of Colorado Boulder for crystallographic data collection. We thank Annette Erbse for her assistance with the resources utilized at the University of Colorado Boulder. We thank Aaron Whiteley, Uday Tak, and Ryan Sayegh for the *Bacillus subtilis PY79* strain and experimental guidance, Amy Palmer for gifting fluorophores, and Lukasz Olenginski and Shea Siwik for critical reading of the manuscript.

## Author contributions

Conceptualization (S.F.S; R.T.B), Investigation (S.F.S.; K.A.D.; R.T.B.), Formal analysis (S.F.S., K.A.D., R.T.B.), Visualization (S.F.S, R.T.B.), Resources (R.T.B.), Writing — Original Draft Preparation (S.F.S., R.T.B.), Writing — Review and Editing (S.F.S., K.A.D., R.T.B.), Funding Acquisition (R.T.B.), Supervision (R.T.B.).

