## Supplementary material for "Structural and functional clues challenge the hypothesis that the *yjdF* riboswitch is natively regulated through broad recognition of azaaromatic compounds": All Supplemental Data

**Table of Contents:**

**Tables**

**Supplemental Table S1**: Sequences of RNAs used in this study.

**Supplemental Table S2**: Crystallographic data and refinement statistics.

**Supplemental Table S3**: Excitation and emission windows used for each fluorogenic ligand.

**Figures**

**Supplemental Figure S1**: Density modified experimental electron density map from iridium-soaked crystal.

**Supplemental Figure S2**: Electron density maps of the native lumichrome-*yjdF* complex.

**Supplemental Figure S3**: Electron density maps of the chelerythrine-*yjdF* complex

**Supplemental Figure S4:** Base-base interactions not predicted from covariation analysis

**Supplemental Figure S5**: Interhelical angles and bends in the *Bsu yjdF* riboswitch.

**Supplemental Figure S6**: Lattice contacts observed around the L4 GAAA tetraloops of the two protomers in the *Bsu yjdF* riboswitch aptamer domain.

**Supplemental Figure S7:** Phylogenetic conservation patterns and in-line probing data superimposed upon the *Bsu yjdF* riboswitch bound to lumichrome.

**Supplemental Figure S8**: Superimposition of the conserved cores of the chelerythrine-bound *Bsu* (blue) and *Rga yjdF* riboswitch (green) aptamer domains.

**Supplemental Figure S9:** Phylogenetic distribution of *yjdF* riboswitches with P2a insertion.

**Supplemental Figure S10**: Secondary structures of select RNAs used in this study.

**Supplemental Figure S11**: Superimposition of T-loop motifs.

**Supplemental Figure S12**: Superimposition of lumichrome- (green) and chelerythrine-bound (orange) *yjdF* aptamer domains.

**Supplemental Figure S13:** Azaaromatic binding cavity in the *Bsu yjdF* riboswitch.

**Supplemental Figure S14**: Superimposition of S-turn motifs

**Supplemental Figure S15**: Magnesium ion bridging phosphate backbone between L2 and J3/4.

**Supplemental Figure S16:** Chelerythrine-induced gene expression of P2a mutations.

**Supplemental Figure S17:** Comparison of the binding pocket of *yjdF*, Peppers fluorogenic aptamer and FMN aptamer.

**Supplemental Table S1**. RNA sequences used in this study.

| **RNA** | **Sequence**^a^ |
| --- | --- |
| *Bacillus subtilis* wild type *yjdF* | GGAUAAAGAAUGAAAAAACACGAUUCGGUUGGUAGUCCGGAUGCAUGAUUGAGAAUGUCAGUAACCUUCCCCUCCUCGGGAUGUCCAUCAUUCUUUA |
| Crystal RNA (6.3.5) | GGAUGAAAAAACACGAUUCGGUUGGUAGUCCGGAUGCCGAAAGGUCAGUAACCUUCCACGAAAGUGGAUGUCCAUCAUCCA |
| L2(GAAA) | GGAUAAAGAAUGAAAAAACACGAUUCGGGAAACCGGAUGCAUGAUUGAGAAUGUCAGUAACCUUCCCCUCCUCGGGAUGUCCAUCAUUCUUUA |
| L3(GAAA) | GGAUAAAGAAUGAAAAAACACGAUUCGGUUGGUAGUCCGGAUGCAUGAAAAUGUCAGUAACCUUCCCCUCCUCGGGAUGUCCAUCAUUCUUUA |
| L4(GAAA) | GGAUAAAGAAUGAAAAAACACGAUUCGGUUGGUAGUCCGGAUGCAUGAUUGAGAAUGUCAGUAACCUUCCCGAAAGGGAUGUCCAUCAUUCUUUA |
| U29C^b^ | GGAUAAAGAAUGAAAAAACACGAUUCGGUUGGUAGCCCGGAUGCAUGAUUGAGAAUGUCAGUAACCUUCCCCUCCUCGGGAUGUCCAUCAUUCUUUA |
| A13G,U29C | GGAUAAAGAAUGAAAAAACGCGAUUCGGUUGGUAGCCCGGAUGCAUGAUUGAGAAUGUCAGUAACCUUCCCCUCCUCGGGAUGUCCAUCAUUCUUUA |
| G24C | GGAUAAAGAAUGAAAAAACACGAUUCGGUUCGUAGUCCGGAUGCAUGAUUGAGAAUGUCAGUAACCUUCCCCUCCUCGGGAUGUCCAUCAUUCUUUA |
| G24C,C73C | GGAUAAAGAAUGAAAAAACACGAUUCGGUUCGUAGUCCGGAUGCAUGAUUGAGAAUGUCAGUAACCUUCCCCUCCUCGGGAUGUCGAUCAUUCUUUA |
| U54A | GGAUAAAGAAUGAAAAAACACGAUUCGGUUGGUAGUCCGGAUGCAUGAUUGAGAAUGUCAGUAACCAUCCCCUCCUCGGGAUGUCCAUCAUUCUUUA |
| C52G | GGAUAAAGAAUGAAAAAACACGAUUCGGUUGGUAGUCCGGAUGCAUGAUUGAGAAUGUCAGUAAGCUUCCCCUCCUCGGGAUGUCCAUCAUUCUUUA |
| U54A,C52G | GGAUAAAGAAUGAAAAAACACGAUUCGGUUGGUAGUCCGGAUGCAUGAUUGAGAAUGUCAGUAAGCAUCCCCUCCUCGGGAUGUCCAUCAUUCUUUA |
| G70C | GGAUAAAGAAUGAAAAAACACGAUUCGGUUGGUAGUCCGGAUGCAUGAUUGAGAAUGUCAGUAACCUUCCCCUCCUCGGGAUCUCCAUCAUUCUUUA |
| C12G | GGAUAAAGAAUGAAAAAAGACGAUUCGGUUGGUAGUCCGGAUGCAUGAUUGAGAAUGUCAGUAACCUUCCCCUCCUCGGGAUGUCCAUCAUUCUUUA |
| C12G,G48C | GGAUAAAGAAUGAAAAAAGACGAUUCGGUUGGUAGUCCGGAUGCAUGAUUGAGAAUGUCACUAACCUUCCCCUCCUCGGGAUGUCCAUCAUUCUUUA |
| A11G | GGAUAAAGAAUGAAAAAGCACGAUUCGGUUGGUAGUCCGGAUGCAUGAUUGAGAAUGUCAGUAACCUUCCCCUCCUCGGGAUGUCCAUCAUUCUUUA |
| A11G, A50G | GGAUAAAGAAUGAAAAAGCACGAUUCGGUUGGUAGUCCGGAUGCAUGAUUGAGAAUGUCAGUGACCUUCCCCUCCUCGGGAUGUCCAUCAUUCUUUA |
| A27G | GGAUAAAGAAUGAAAAAACACGAUUCGGUUGGUGGUCCGGAUGCAUGAUUGAGAAUGUCAGUAACCUUCCCCUCCUCGGGAUGUCCAUCAUUCUUUA |
| A27G, U49C | GGAUAAAGAAUGAAAAAACACGAUUCGGUUGGUGGUCCGGAUGCAUGAUUGAGAAUGUCAGCAACCUUCCCCUCCUCGGGAUGUCCAUCAUUCUUUA |
| A27G, U49C, U22C | GGAUAAAGAAUGAAAAAACACGAUUCGGCUGGUGGUCCGGAUGCAUGAUUGAGAAUGUCAGCAACCUUCCCCUCCUCGGGAUGUCCAUCAUUCUUUA |
| U23A | GGAUAAAGAAUGAAAAAACACGAUUCGGUAGGUAGUCCGGAUGCAUGAUUGAGAAUGUCAGUAACCUUCCCCUCCUCGGGAUGUCCAUCAUUCUUUA |
| U23G | GGAUAAAGAAUGAAAAAACACGAUUCGGUGGGUAGUCCGGAUGCAUGAUUGAGAAUGUCAGUAACCUUCCCCUCCUCGGGAUGUCCAUCAUUCUUUA |
| C53G,G70C | GGAUAAAGAAUGAAAAAACACGAUUCGGUUGGUAGUCCGGAUGCAUGAUUGAGAAUGUCAGUAACGUUCCCCUCCUCGGGAUCUCCAUCAUUCUUUA |
| C53U,G70A | GGAUAAAGAAUGAAAAAACACGAUUCGGUUGGUAGUCCGGAUGCAUGAUUGAGAAUGUCAGUAACUUUCCCCUCCUCGGGAUAUCCAUCAUUCUUUA |
| C52U,U71C | GGAUAAAGAAUGAAAAAACACGAUUCGGUUGGUAGUCCGGAUGCAUGAUUGAGAAUGUCAGUAAUCUUCCCCUCCUCGGGAUGCCCAUCAUUCUUUA |
| +P2a | GGAUAAAGAAUGAAAAAACACAACGGCGAAAGCCAGAUUCGGUUGGUAGUCCGGACGCAUGAUUGAGAAUGUCAGUAACCUUCCCCUCCUCGGGAUGUCCAUCAUUCUUUA |
| +P2a/U23A | GGAUAAAGAAUGAAAAAACACAACGGCGAAAGCCAGAUUCGGUAGGUAGUCCGGACGCAUGAUUGAGAAUGUCAGUAACCUUCCCCUCCUCGGGAUGUCCAUCAUUCUUUA |

^a^color scheme: yellow, P1 helix; green, P2 helix; magenta, P3 helix; cyan, P4 helix; gray, terminal loops L2, L3 and L4; blue, P2a insertion; red, mutation

^b^number scheme used for mutants is that of the crystallized RNA and the resultant models deposited in the PDB

**Supplemental Table S2.** Crystallographic Table of Data and Refinement Statistics.

| Crystal | *Bsu-yjdF,* iridium  (lumichrome) | *Bsu-yjdF,* native  (lumichrome) | *Bsu-yjdF,* native  (chelerythrine) |
| --- | --- | --- | --- |
| PDB ID | 9EBP | 9EBV | 9EC4 |
| **Data collection** |  |  |  |
| Wavelength (Å) | 1.1055 | 1.5418 | 1.5418 |
| Resolution Range (Å)^a^ | 46 – 2.7 (2.8 – 2.7) | 20 – 2.52 (2.59 – 2.52) | 20 – 2.85 (2.95 – 2.85) |
| Space Group | P2_1_2_1_2 | P2_1_2_1_2 | P2_1_2_1_2 |
| Unit Cell |  |  |  |
| a, b, c (Å) | 111.59, 65.84, 81.25 | 110.72, 68.93, 82.65 | 109.65, 69.27, 82.10 |
| α, β, γ (°) | 90, 90, 90 | 90, 90, 90 | 90, 90, 90 |
| Unique Reflections | 16938 (1594) | 21609 (1879) | 14791 |
| R_merge_ | 0.091 (1.21) | 0.074 (0.545) | 0.045 (0.535) |
| R_pim_ | 0.025 (0.35) | 0.025 (0.322) | 0.026 (0.313) |
| Redundancy | 13.7 (2.4) | 4.8 (3.4) | 2.3 (2.0) |
| Completeness (%) | 99.7 (99.0) | 97.4 (77.6) | 94.9 (90.0) |
| I/σ(I) | 20.4 (1.5) | 20.6 (1.6) | 17.8 (2.0) |
| CC_1/2_ | 1.00 (0.815) | --- (0.854) | --- (0.923) |
| **Refinement** |  |  |  |
| Resolution (Å) | 46 – 2.7 (2.8 – 2.7) | 20 – 2.52 (2.61 -2.52) | 20 – 2.85 (2.95 – 2.85) |
| No. reflections | 16899 (1594) | 21432 (1879) | 14481 (1535) |
| R_work_/R_free_ | 0.25/0.29 (0.46/0.48) | 0.22/0.25 (0.32/0.36) | 0.23/0.26 (0.38/0.43) |
| No. atoms |  |  |  |
| Total | 3603 | 3600 | 3638 |
| RNA | 3420 | 3420 | 3420 |
| Ligands | 162 | 109 | 130 |
| Water | 21 | 71 | 88 |
| B-factors |  |  |  |
| Total | 62.8 | 56.4 | 58.8 |
| RNA | 62.1 | 56.2 | 58.2 |
| Ligands | 77.5 | 65.2 | 70.7 |
| Water | 63.9 | 54.4 | 64.9 |
| r.m.s. deviations |  |  |  |
| Bond Length (Å) | 0.013 | 0.012 | 0.012 |
| Bond Angle (°) | 1.40 | 1.07 | 0.98 |

**Supplemental Table S3.** Excitation and emission wavelengths used for fluorescence measurements.

| Fluorogen | Excitation (nm)^a^ | Emission (nm)^a^ | Binding (if different)^b^ |
| --- | --- | --- | --- |
| Thiazole Orange | 499-12 | 536-12 |  |
| Chelerythrine | 330-10 | 550-10 | 330-20/570-60 |
| Proflavine | 464-8 | 502-16 |  |
| DAPI | 370-10 | 460-10 |  |
| Malachite Green | 617-10 | 653-10 |  |
| Thioflavin T | 446-10 | 483-10 |  |
| Biliverdin | 448-10 | 490-10 |  |
| HBC620 | 577-10 | 620-10 |  |
| NMMP | 400-8 | 610-10 |  |
| DFHO | 505-10 | 545-10 |  |

^a^Window is given as x-y, where x is the wavelength and y is the bandwidth.

^b^Binding measurements to determine ligand affinity given as excitation/emission.

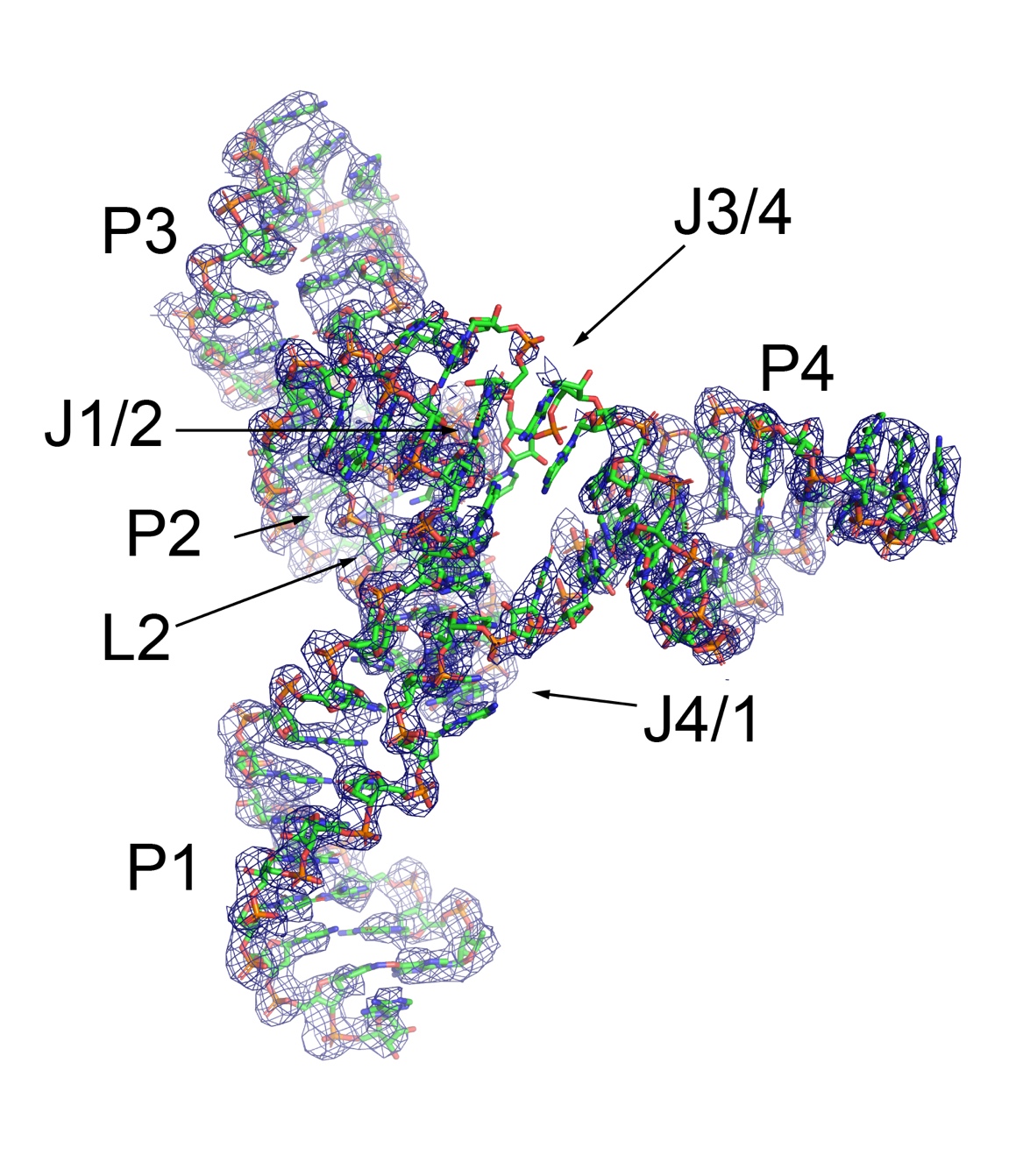

**Supplemental Figure S1.** Density modified experimental electron density map from iridium-soaked crystal. The molecule shown is protomer A of the final model built from the iridium dataset (PDB 9EBP). The electron density map is contoured at 1.0 sigma and within 2.0 Å of model atoms. Note that the paired helices (P1-P4), terminal loops, J1/2 and J3/4 are well defined by the density while J3/4 is poorly defined in the initial map.

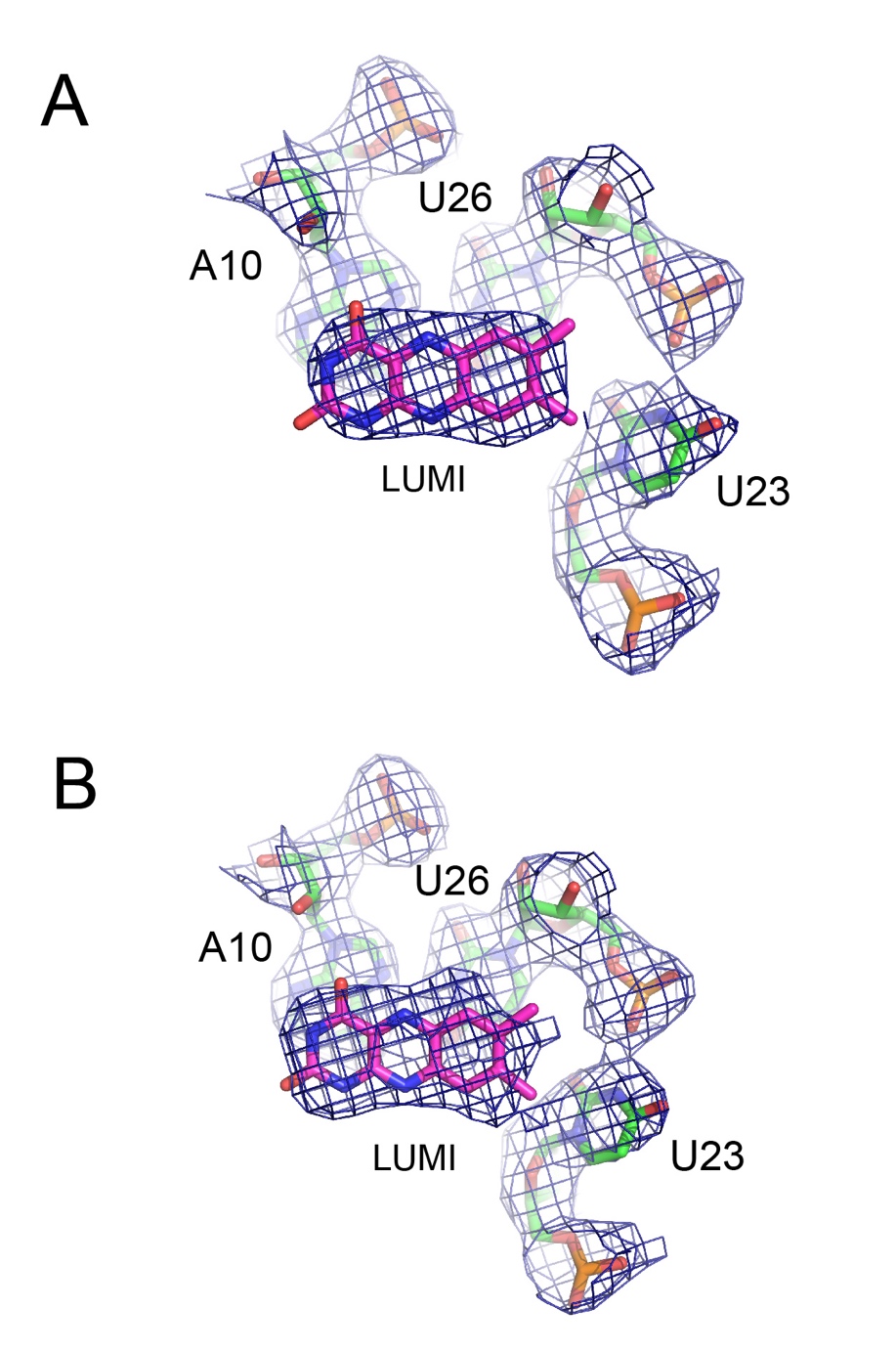

**Supplemental Figure S2.** Electron density maps of the native lumichrome-*yjdF* complex (PDB 9EBV). (A) 2Fo-Fc electron density map of ligand binding core including nucleotides A10, U23, and U26 (green) and lumichrome (magenta), contoured at 1.2 sigma and within 2.0 Å of model atoms. (B) Composite omit map of ligand binding core (same perspective as in panel (A)), contoured at 1.2 sigma and within 2.0 Å of model. The 2Fo-Fc map suggests some orientational ambiguity of lumichrome, but the pose in the final model was based upon that picked automatically by LigandFit in PHENIX (independently in both protomers) and the observation that in the other pose (with the carbonyl groups adjacent to U23), there would be a significant electrostatic clash between a lumichrome carbonyl oxygen and the bridging phosphate oxygen of U26.

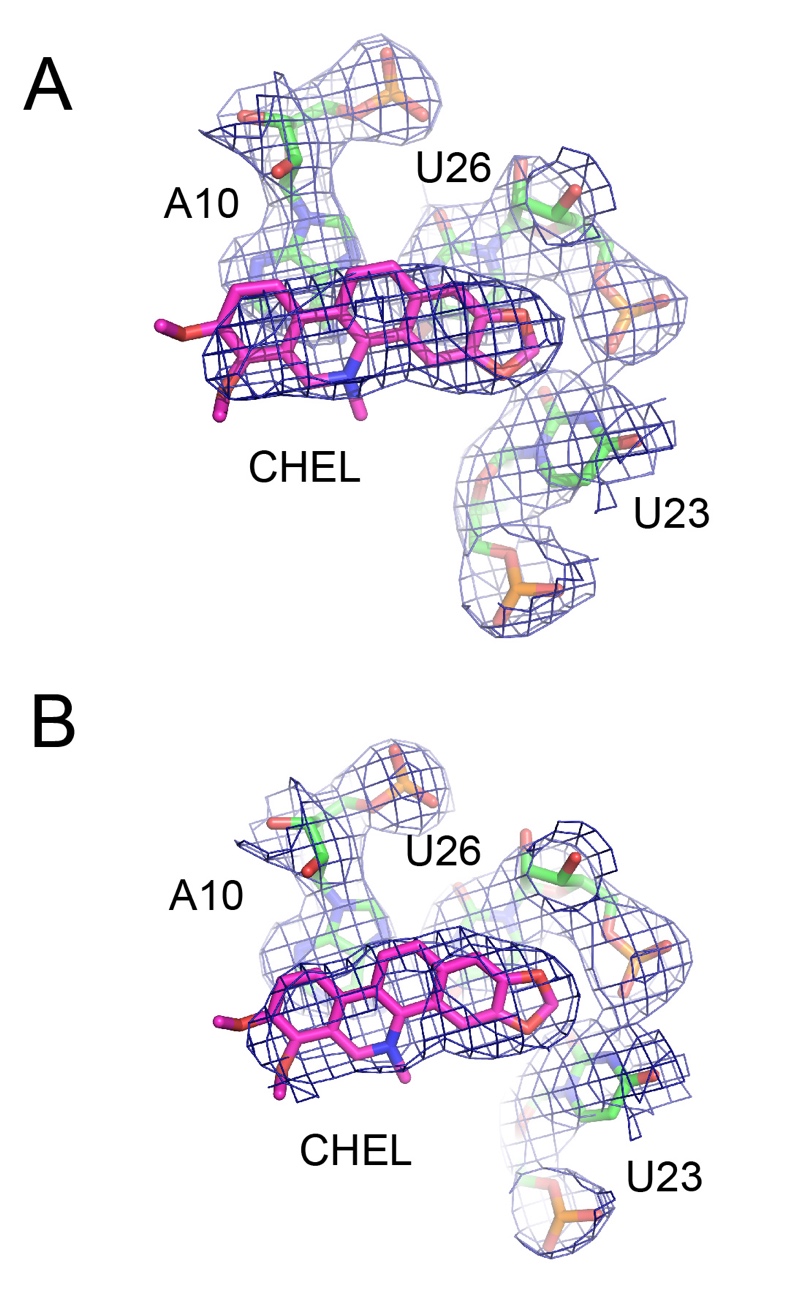

**Supplemental Figure S3.** Electron density maps of the chelerythrine-*yjdF* complex (PDB 9EC4). (A) 2Fo-Fc electron density map of ligand binding core including nucleotides A10, U23, and U26 (green) and chelerythrine (magenta), contoured at 1.2 sigma and within 2.0 Å of model atoms. (B) Composite omit map of ligand binding core (same perspective as in panel (A)), contoured at 1.2 sigma and within 2.0 Å of model. In both cases, the density for the ligand is well defined except for the methyl groups of the methoxy moieties.

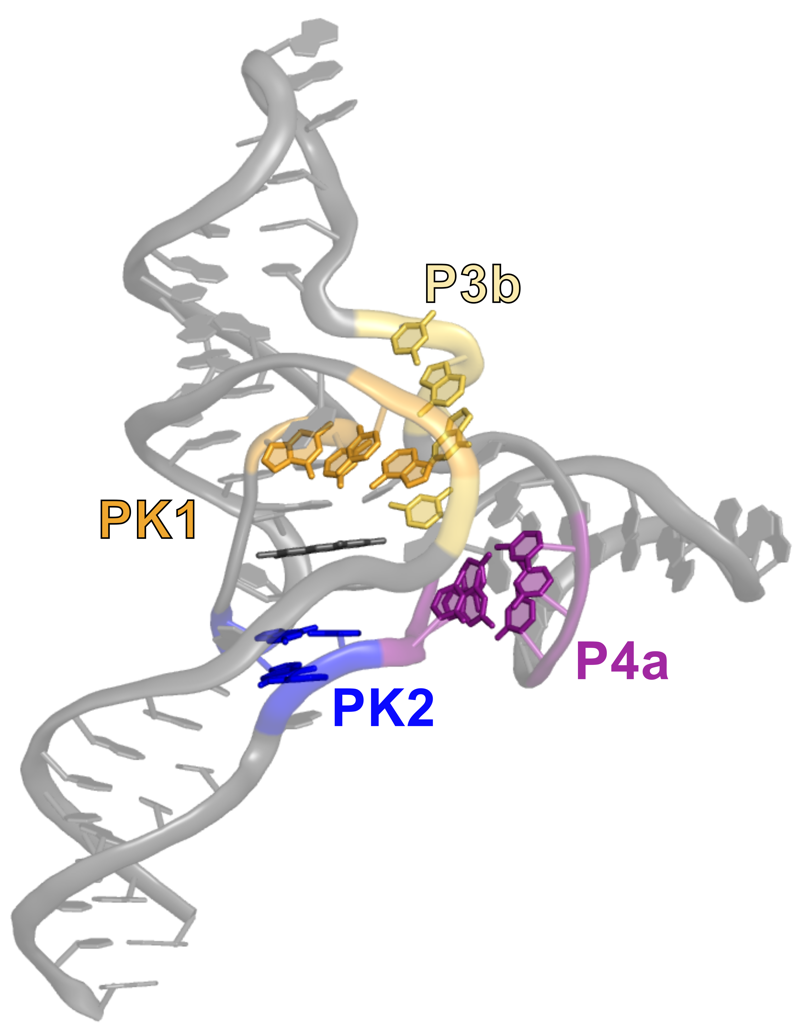

**Supplemental Figure S4.** Global structure of the *yjdF* riboswitch aptamer domain with base pairs that were not predicted from covariation analysis of phylogenetic variants emphasized. Base pairs in PK1 (orange), PK2 (blue), P3b (yellow-gold), and P4a (purple) are colored while the predicted base pairs of helices P1-P4 and the engineered terminal loops are in grey. Note that these base pairs surround the ligand binding pocket (PK1, PK2) or part of a potential communication relay between the ligand binding pocket and the anti-RBS in L4 (P3b and P4a).

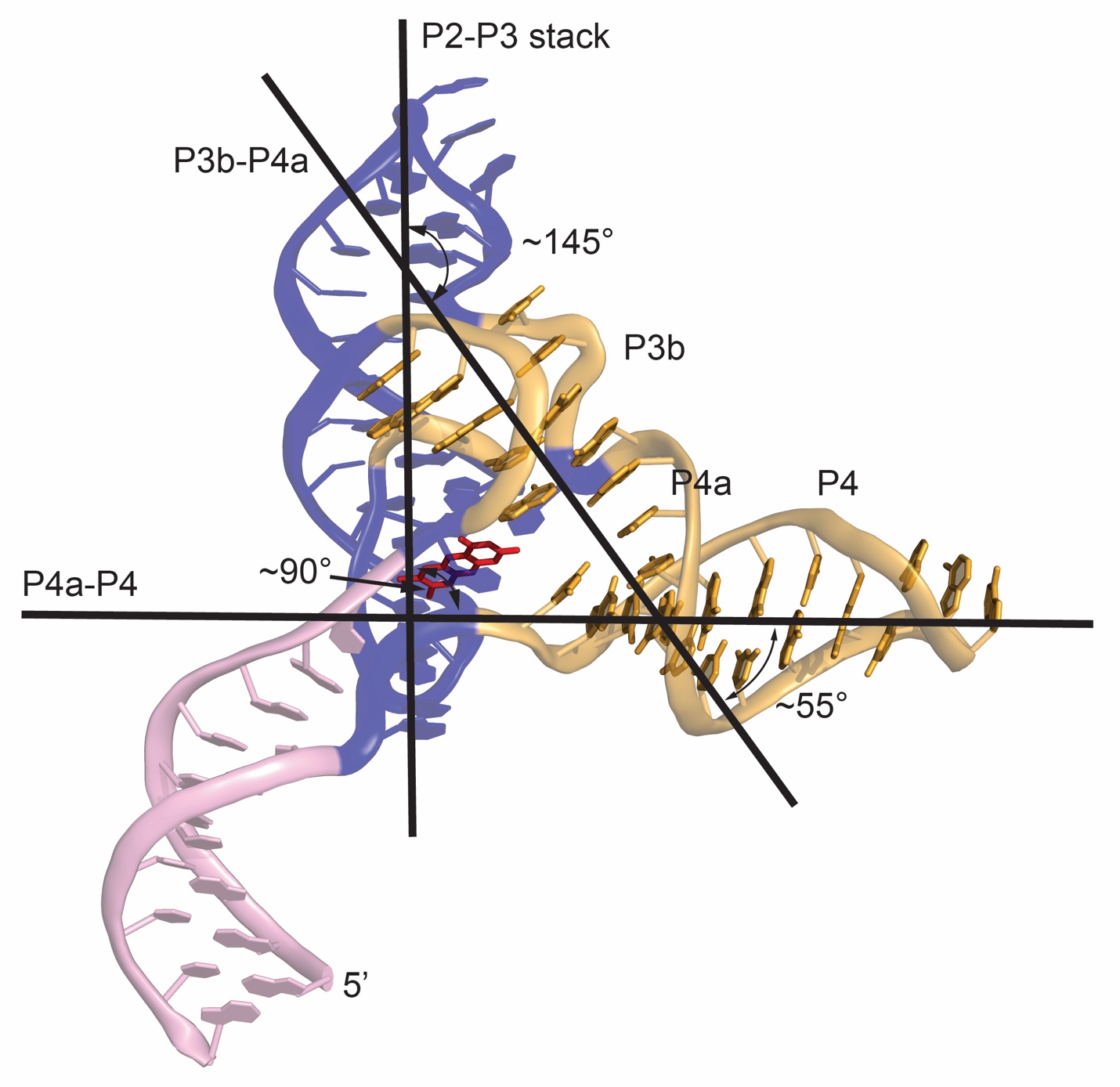

**Supplemental Figure S5.** Interhelical angles and bends in the *Bsu yjdF* riboswitch.

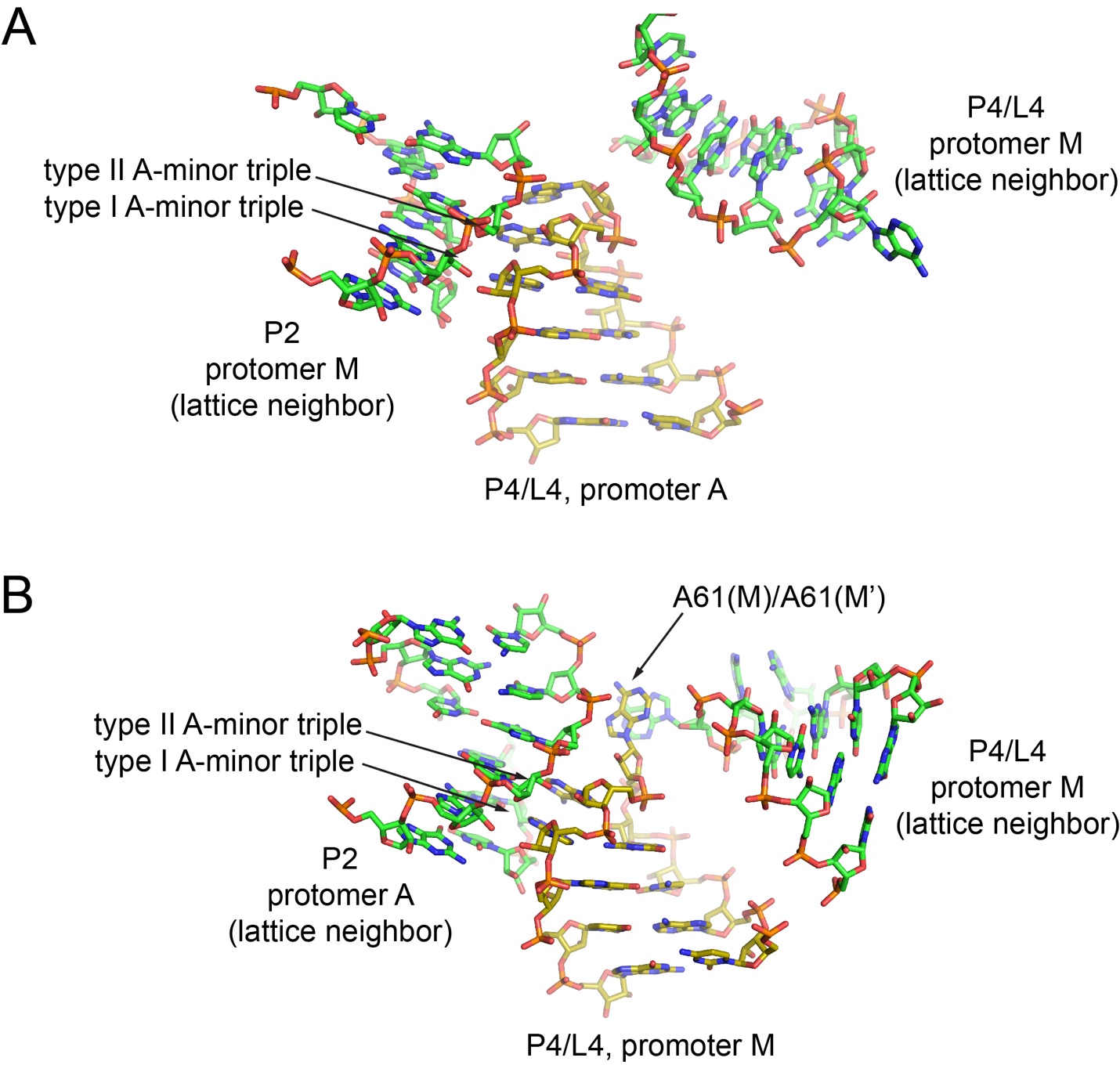

**Supplemental Figure S6.** Lattice contacts observed around the L4 GAAA tetraloops of the two protomers in the *Bsu yjdF* riboswitch aptamer domain. (A) L4 GAAA tetraloop of protomer A primarily contacts the P2 stem via standard A-minor triple interactions using the most common conformation of the GAAA tetraloop in which all three adenosine nucleobases are stacked. (B) L4 GAAA tetraloop of protomer M contacts the P2 stem of a neighboring molecule using standard A-minor triple interactions, but the adenosine at position 2 of the tetraloop is destacked from the adenosine at position 3 and forms a stacking interaction with the second position adenosine of a neighboring protomer M tetraloop.

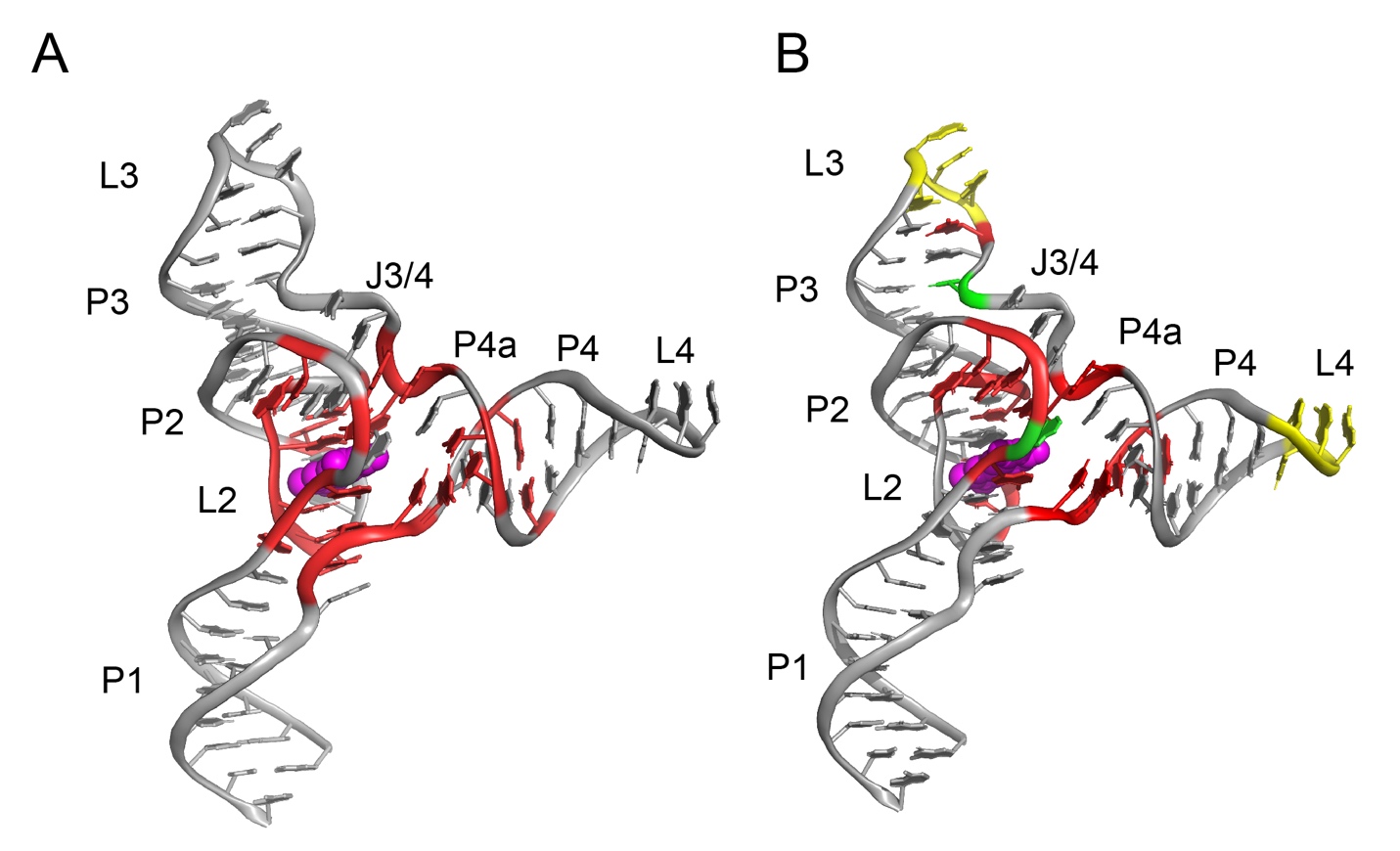

**Supplemental Figure S7.** (A) Nucleotides whose identity is >98% conserved across *yjdF* riboswitch phylogenetic variants are colored in red while the rest of the aptamer domain is colored in grey and the ligand in magenta. (B) Nucleotides that display “in-line” cleavage protection (red), enhancement (green), or no change (yellow) between the unbound and chelerythrine-bound states are displayed on the structure of the *yjdF* riboswitch. Grey nucleotides showed no cleavage in either the apo or bound states. Note that the no change nucleotides were observed in the wild-type *B. subtilis* structure, but we have colored the GAAA tetraloops of the crystallized variant to highlight their analogous positions.

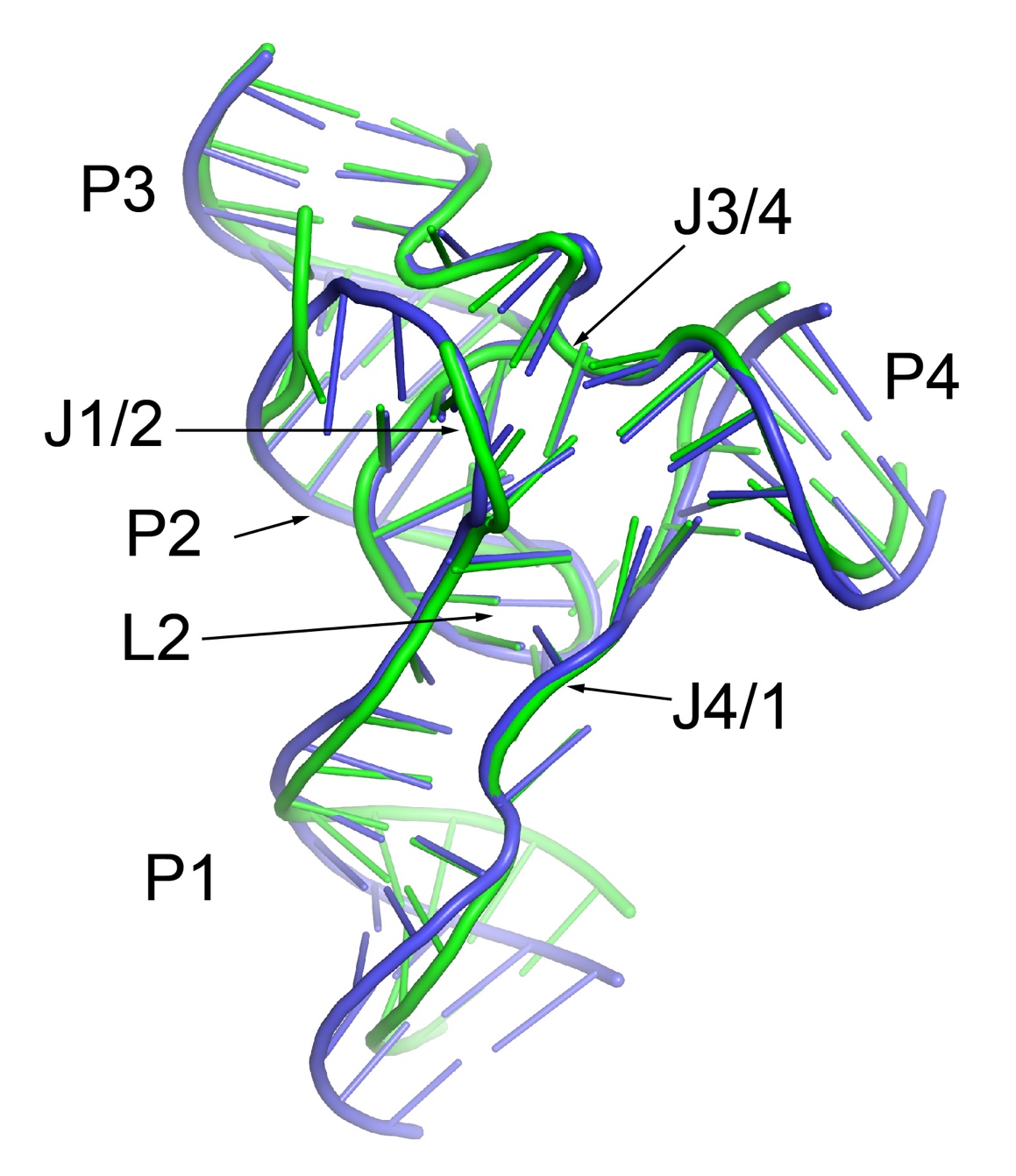

**Supplemental Figure S8.** Superimposition of the conserved cores of the chelerythrine-bound *Bsu* (blue) and *Rga yjdF* riboswitch (green) aptamer domains. For superimposition, nucleotides 5-17, 44-67, 90-104, and 111-125 from protomer N of the *Rga yjdF* RNA (PDB 8UIW) and nucleotides 1-38, 43-57, and 66-80 from protomer M of the *Bsu yjdF* RNA (PDB 9EC4) were used. Superposition of the phosphate, nonbridging phosphate oxygens, and ribose carbons from each RNA were superimposed using the align command in PyMOL. The resulting rmsd over 393 atoms in each RNA was 1.37 Å. RNAs are represented in cartoon format.

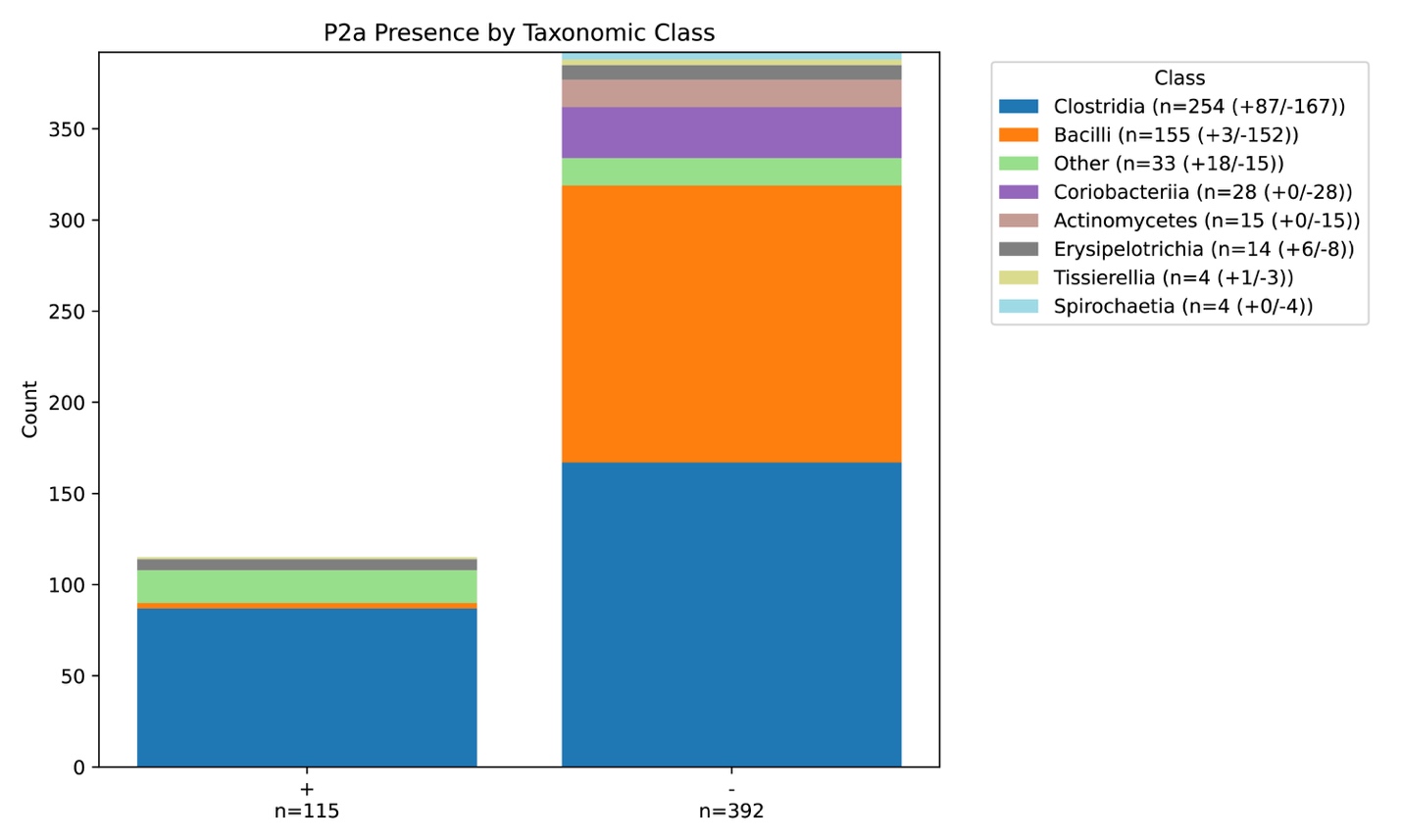

**Supplemental Figure S9.** Distribution of *yjdF* riboswitches with (+, left) and without (-, right) the P2a insertion. All bacterial classes with a single *yjdF* variant (singletons) were grouped into the “other” class. In the legend, the number of variants with and without P2a are denoted for each class.

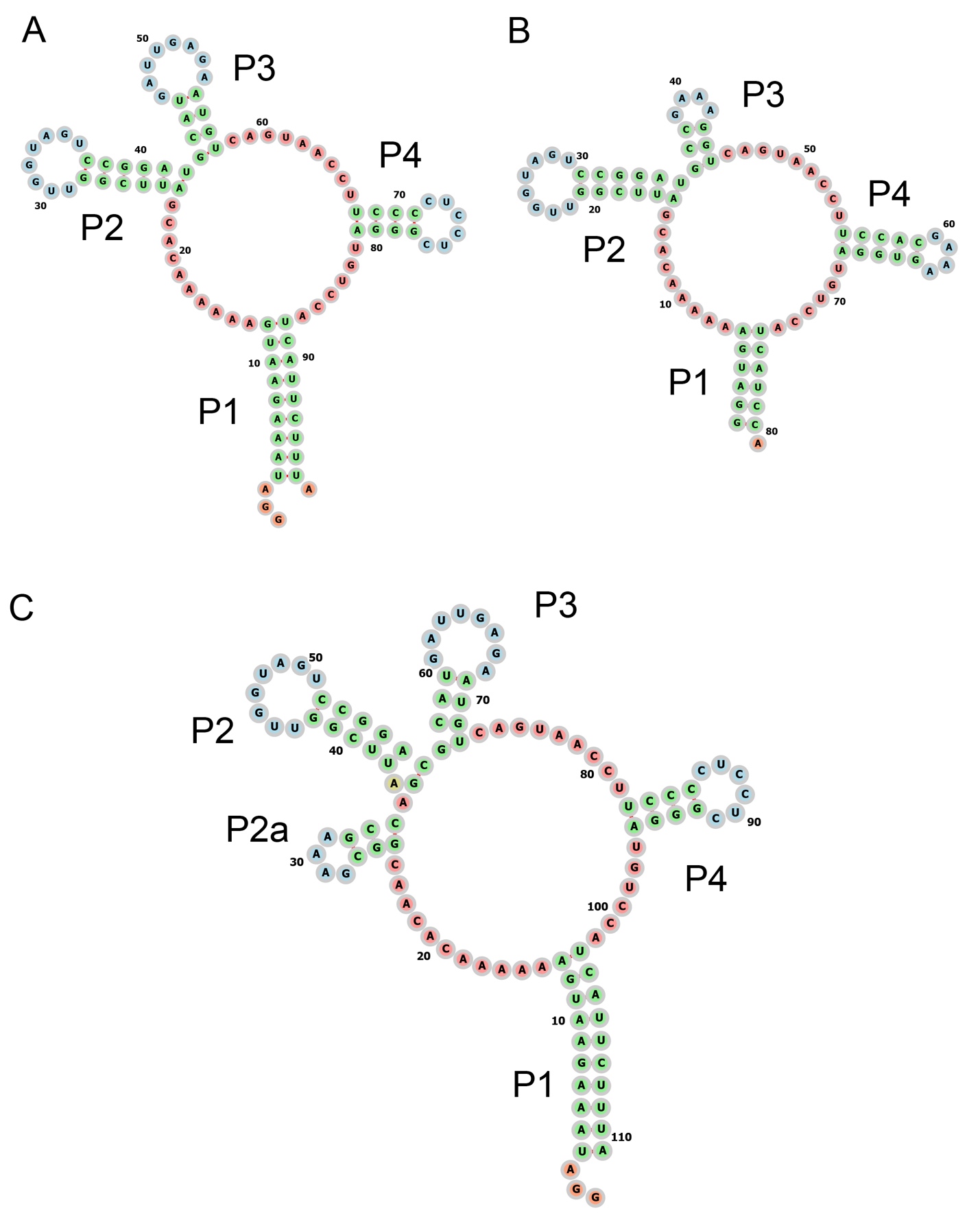

**Supplemental Figure S10.** Secondary structures of the (A) wild type *Bsu* *yjdF* aptamer domain (B) *Bsu* *yjdF* RNA used for crystallization and (C) *Bsu yjdF* with the P2a insertion.

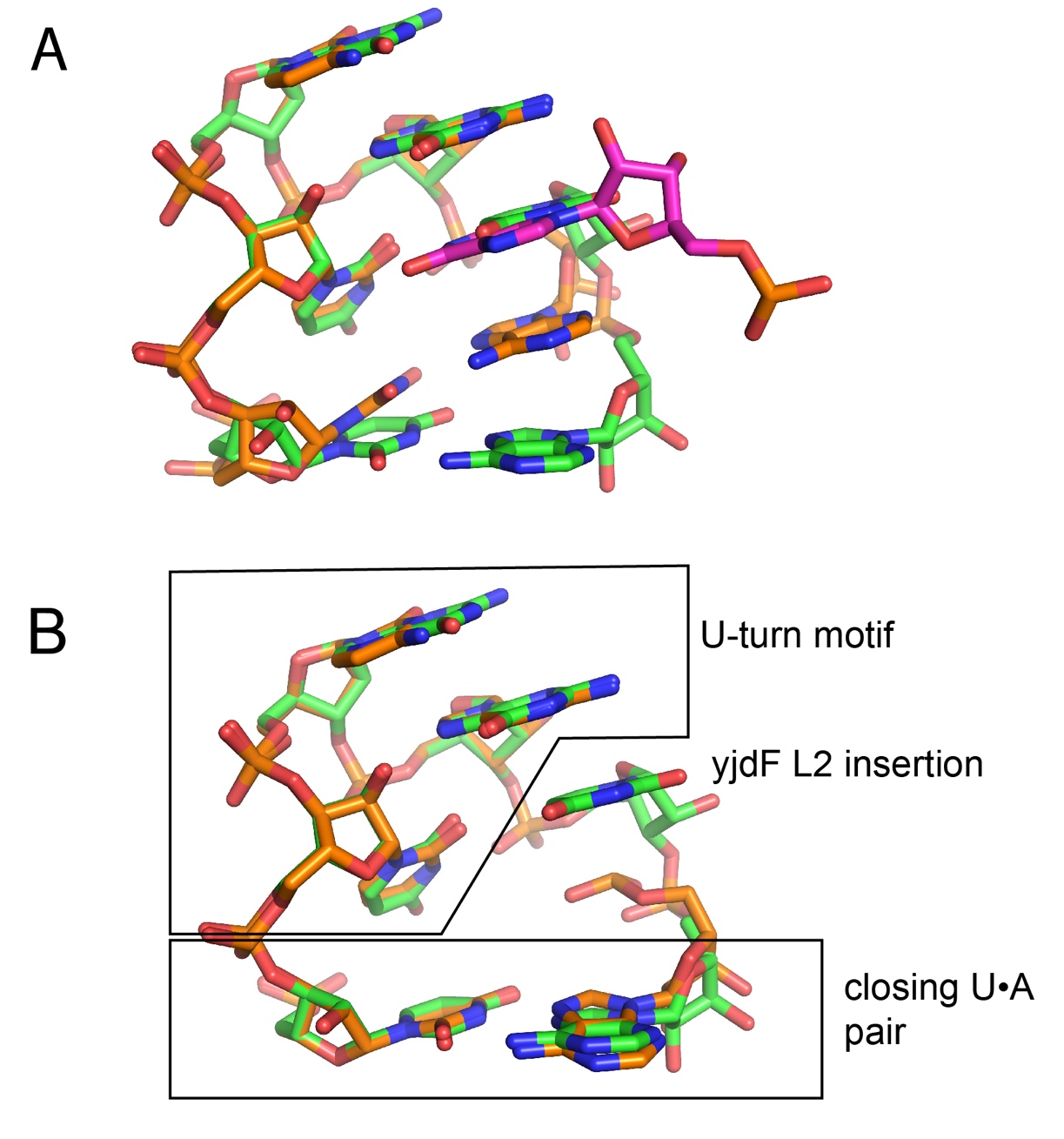

**Supplemental Figure S11.** Superimposition of T-loops. (A) Unbiased superimposition of the full T-loop motif of *yjdF* (nucleotides 22-27, green) with the T-loop motif of unmodified tRNA^Asp^ (PDB ID 6UGG; nucleotides 54-58, orange) using the align command in PyMOL. Note that the U-turn motif superimposes very well and that the fourth nucleotide of the L2 T-loop motif is placed where the D-loop adenosine (magenta) docks into the tRNA T-loop. (B) Superimposition of fragments of the tRNA T-loop motif onto *yjdF* L2 as two separate fragments: the U-turn motif (nucleotides 55-57) and the closing U•A pair (nucleotides 54 and 58). These two fragments superimpose well and clearly reveal that the fourth nucleotide of the *yjdF* L2 (nucleotide 26) represents the insertion element.

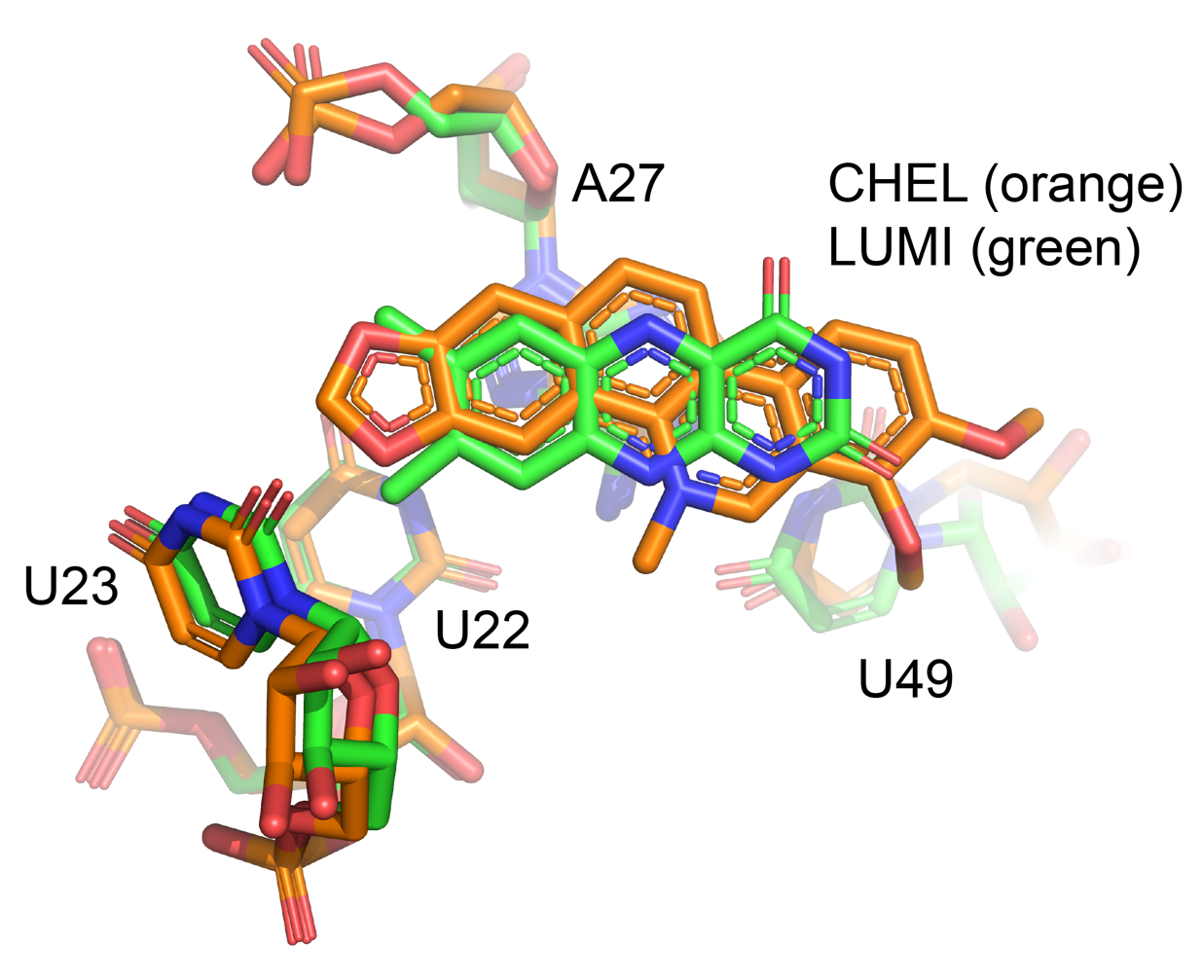

**Supplemental Figure S12.** Superimposition of lumichrome- (green) and chelerythrine-bound (orange) *yjdF* aptamer domains. Superimposition of both structures was performed using U22, U23, A27, and U49 yielding an RMSD = 0.47 Å. In both structures, the ligand is centered above A27 of the U22•A27-U49 base triple and in Van der Waals contact with U23 of the L2 loop.

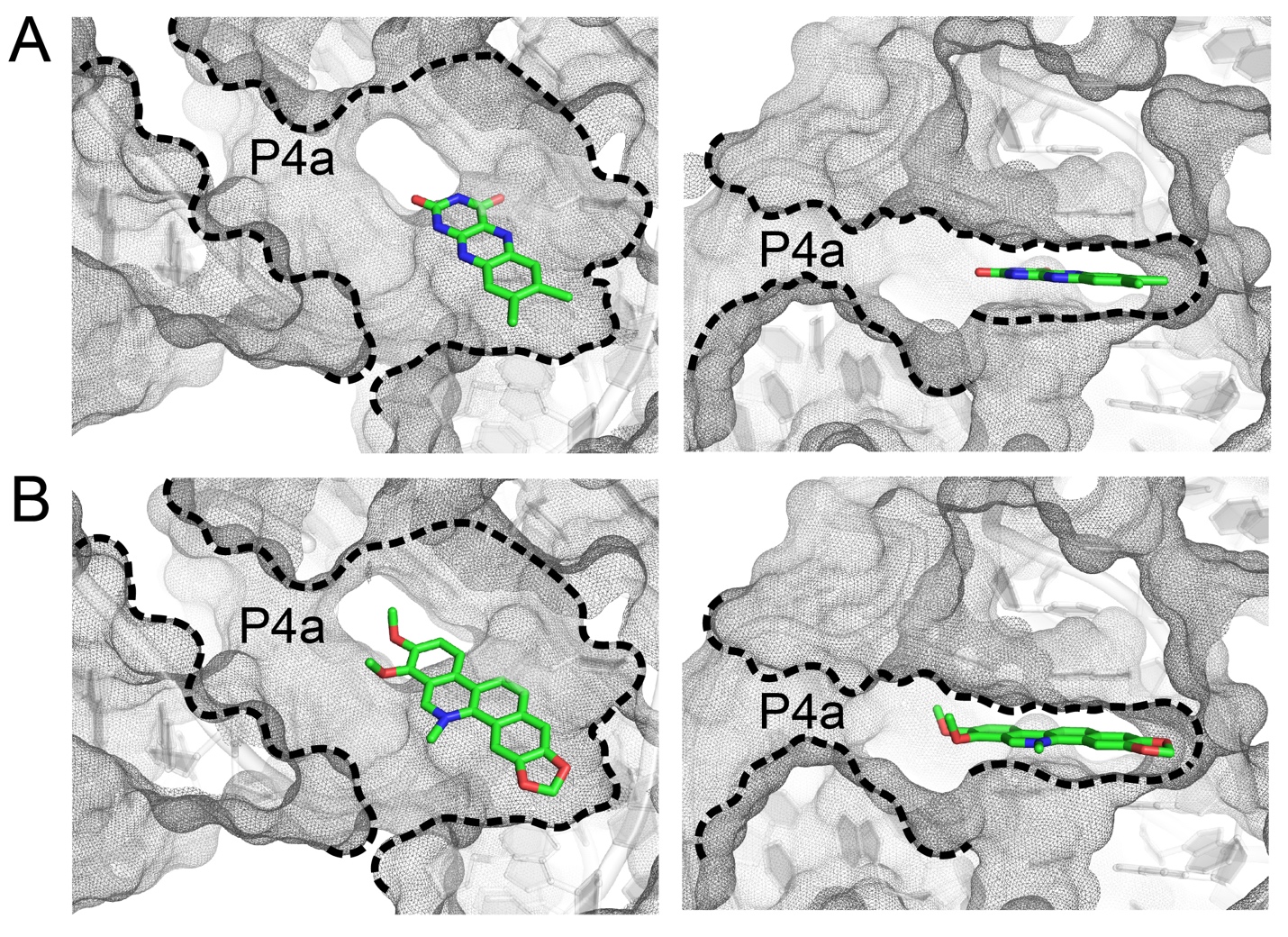

**Supplementary Figure S13.** Cross section through the ligand binding pocket in the B. subtilis *yjdF* riboswitch bound to (A) lumichrome and (B) chelerythrine. The open pocket is continuous with the major groove of P4a (left views) that is distinct from the flat “aromatic” binding region (right views). The pocket is also open to bulk solvent towards the “bottom” of the pocket in the left views. In the view from other perspectives, this opening is much wider and able to accommodate large non-aromatic substituents such as that of staurosporine.

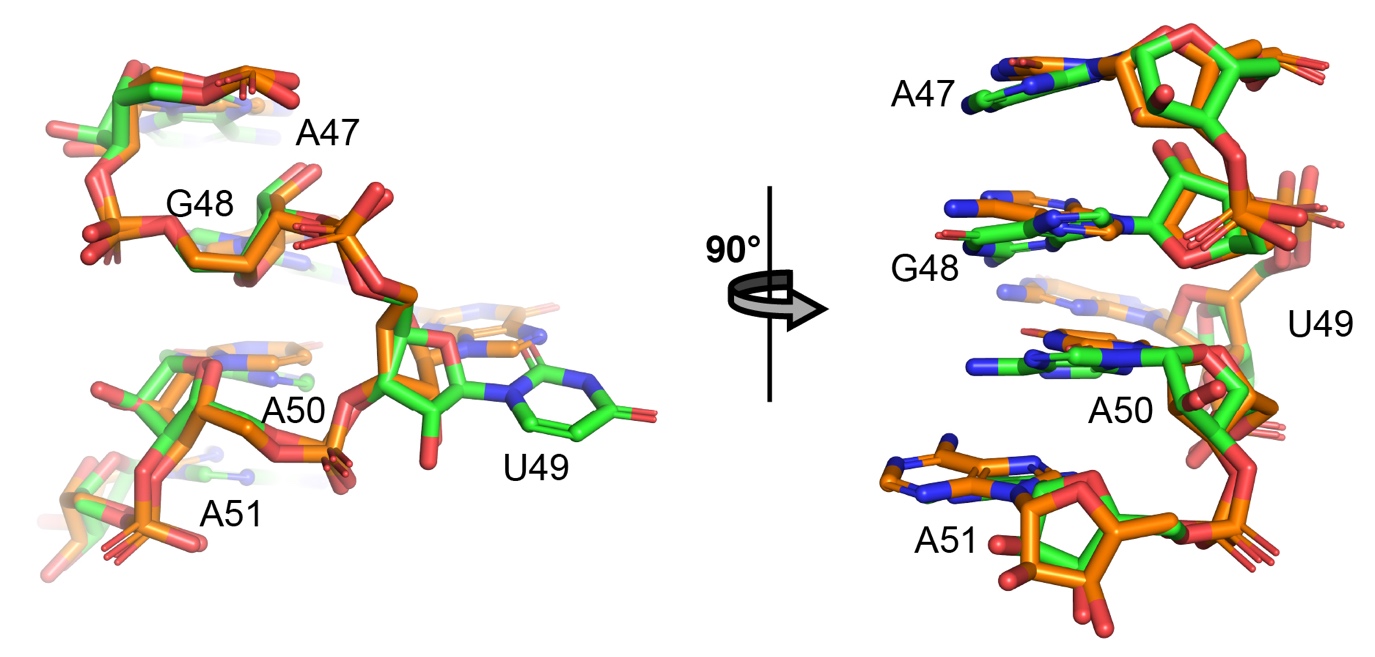

**Supplemental Figure S14.** The S-turn motif in J3/4 of *yjdF* (nucleotides 47-51, green) superimposed upon the S-turn motif of the T-box motif (nucleotides 65-69, chain B, PDB ID 6UFM, orange). Superimposition results in an RMSD of 0.63 Å over backbone atoms. Numbering in the figure corresponds to the *yjdF* riboswitch.

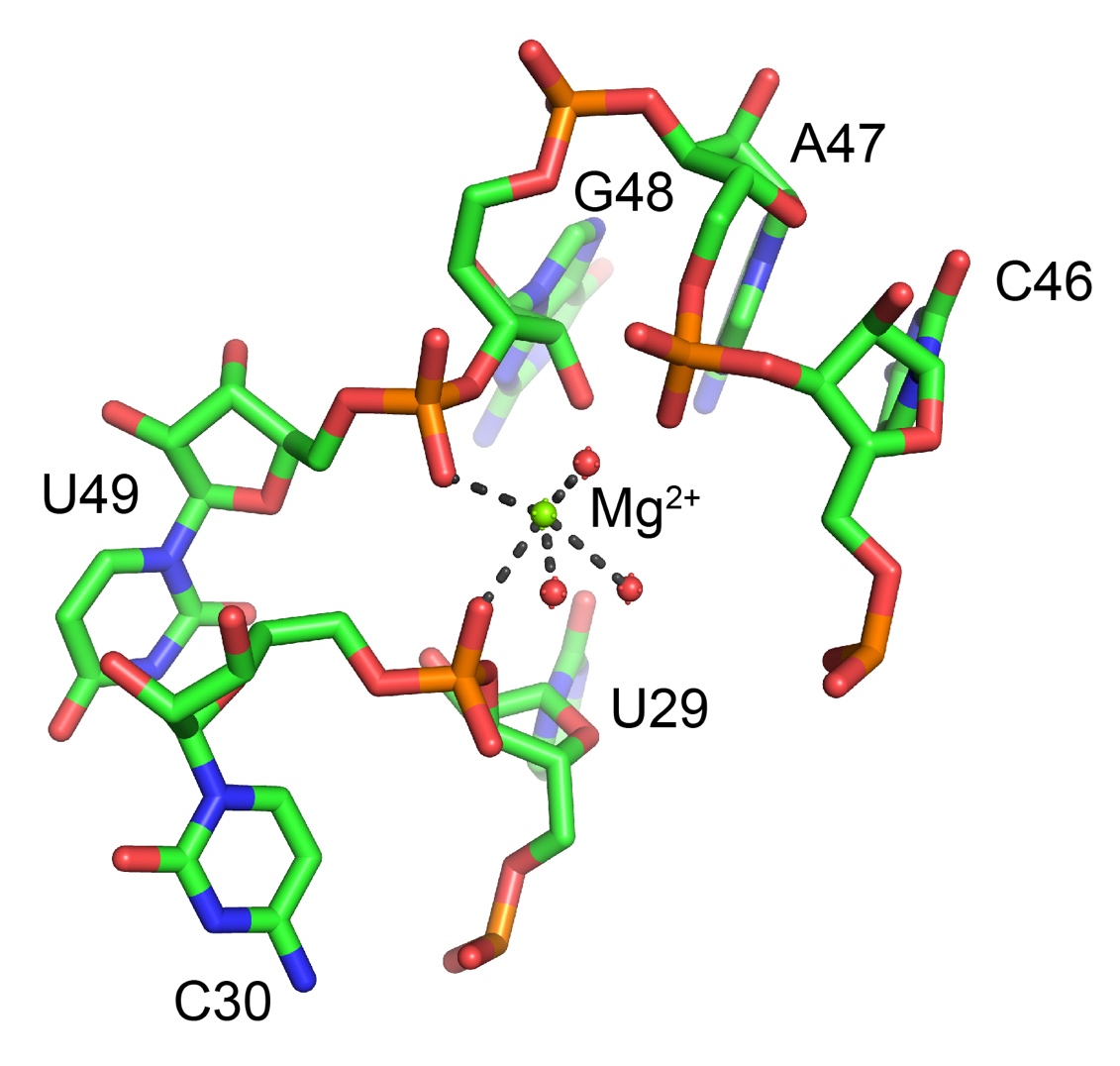

**Supplemental Figure S15.** Magnesium that forms two inner sphere coordinations with non-bridging phosphate oxygens (C30 and U49).The sharp bend in the helix between U29 and C30 at the junction between the 3′-end of L2 and the beginning of the 3′-side of P2 while U49 is in the S-turn in J3/4 and forms part of a triple pair that is adjacent to the aromatic ligand.

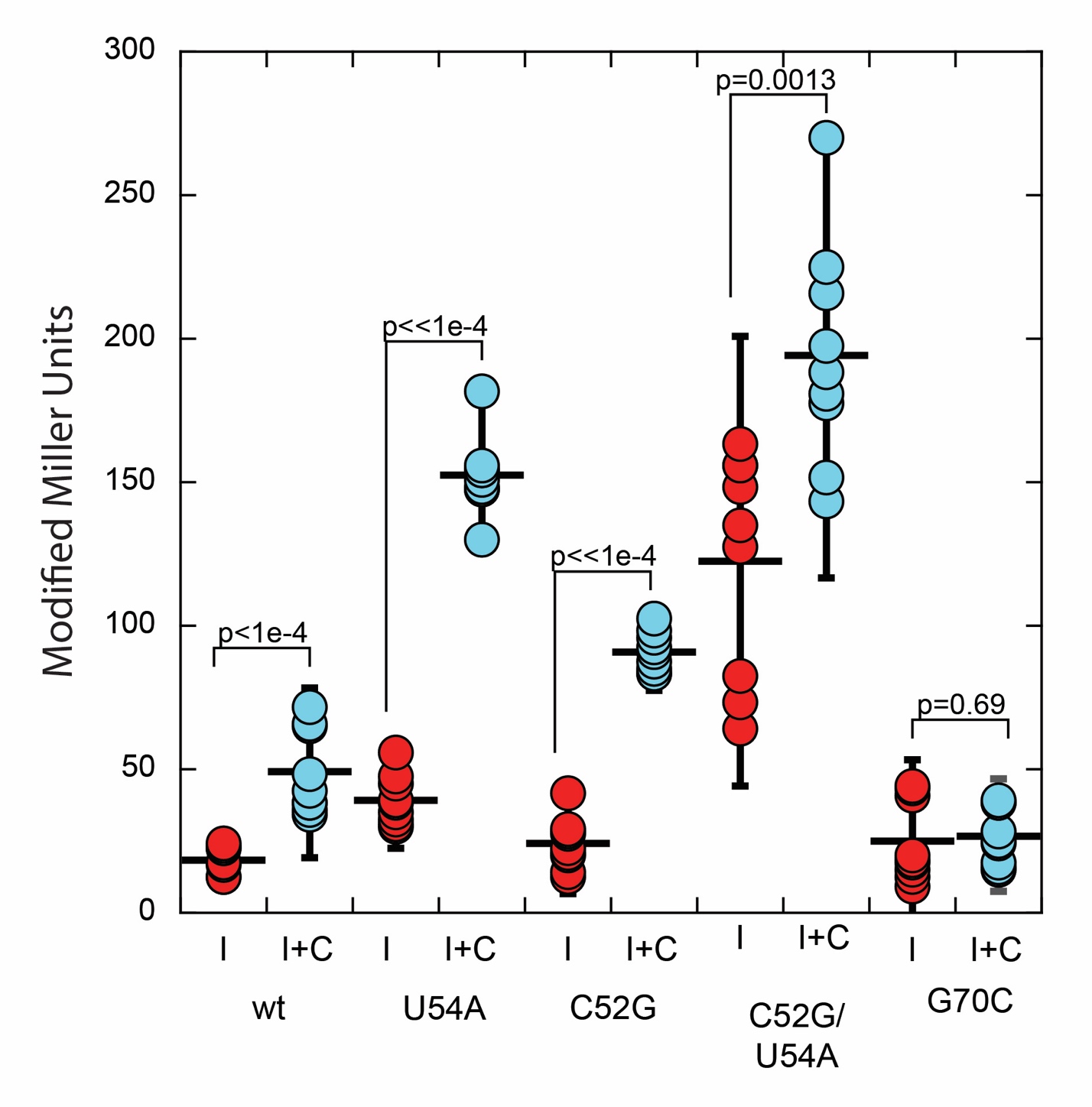

**Supplemental Figure S16.** Miller assay results of mutations in P4a designed to either strengthen (U54A and C52G) or weaken (G70C) the P4a helix in the absence (I, IPTG only) or presence of 50 nM chelerythrine (I+C). While the U54A and C52G support nearly wild-type regulatory activity, C52G/U54A displays only a modest increase in reporter expression in the presence of chelerythrine, while G70C is not able to activate gene expression. The error bars correspond to two standard deviations from the mean and the p-values were calculated using a student’s two-tailed t-test using Excel.

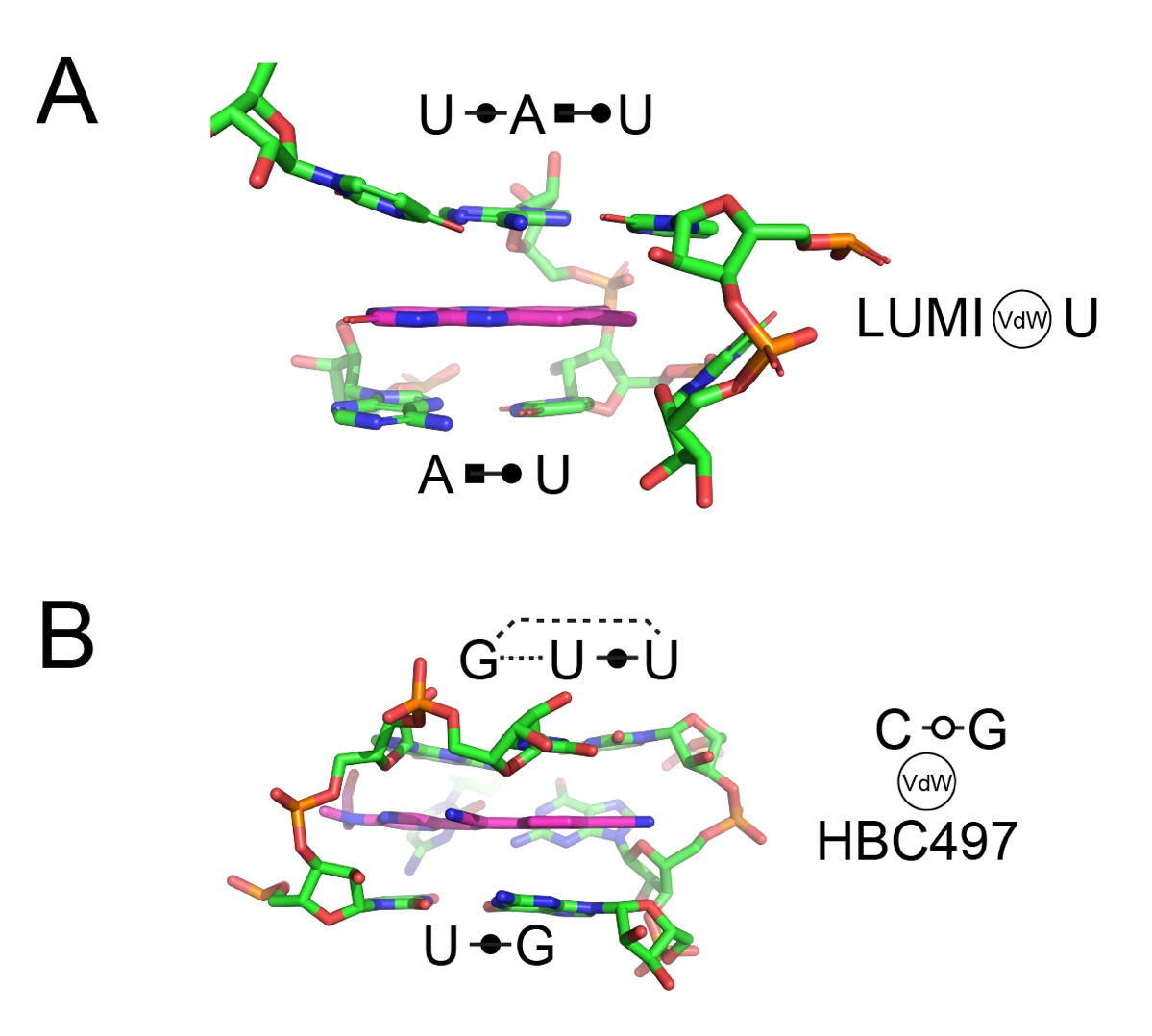

**Supplemental Figure 17.** Bases and base-base interactions that create large, aromatic binding pockets in the (A) *B. subtilis yjdF* riboswitch and (B) Peppers fluorogenic aptamer (PDB ID 7EOL). In each case, the pocket is created by a base triple (top in each view) and a non-WCF base pair on the bottom. Each pocket also is flanked by an inclined single base (*yjdF*) or base pair (Peppers) that form direct interactions with the edge of the aromatic ligand.
